# Defining relative mutational difficulty to understand cancer formation and prevention

**DOI:** 10.1101/789313

**Authors:** Lin Shan, Jiao Yu, Zhengjin He, Shishuang Chen, Mingxian Liu, Hongyu Ding, Liang Xu, Jie Zhao, Ailing Yang, Hai Jiang

## Abstract

Most mutations in human cancer are low-frequency missense mutations, whose functional status remains hard to predict. Here we show that depending on the type of nucleotide change and the surrounding sequences, the tendency to generate each type of nucleotide mutations varies greatly, even by several hundred folds. Therefore, a cancer-promoting mutation may appear only in a small number of cancer cases, if the underlying nucleotide change is too difficult to generate. We propose a method that integrates both the original mutation counts and their relative mutational difficulty. Using this method, we can accurately predict the functionality of hundreds of low-frequency missense mutations in p53, PTEN and INK4A. Many loss-of-function p53 mutations with dominant negative effects were identified, and the functional importance of several regions in p53 structure were highlighted by this analysis. Furthermore, mutational difficulty analysis also points to potential means of cancer prevention. Our study not only established relative mutational difficulties for different types of mutations in human cancer, but also showed that by incorporating such parameter, we can bring new angles to understanding cancer formation and prevention.

## Introduction

Gene mutation is a major cause of tumorigenesis. Certain mutations on important cancer genes such as KRAS and p53 drive cancer formation(Cheok et al., 2011; Kastenhuber and Lowe, 2017). As a result, such mutations are enriched in cancer, and are found in numerous cancer samples. It is generally perceived that if a mutation occurs in higher number of cancer cases, it is more likely to be a driver mutation(Baugh et al., 2018; Brosh and Rotter, 2009). However, the vast majority of mutations in cancer only occurs in very small number of cancer cases, and the functional impacts of these mutations are hard to predict.

To address this problem, it is necessary to consider that, the chance of observing a mutation in cancer cases is influenced by at least two major factors: 1) how difficult it is to generate the mutation; and 2) whether the mutation promotes cancer, therefore it will be selectively enriched in cancer cases. If different mutations are initially generated at significantly different rates, it will greatly impact the mutational distribution in cancer genome database such as COSMIC (Catalogue Of Somatic Mutations In Cancer). Certain cancer-driving, but too-hard-to generate mutations may appear exceedingly rare in cancer database, yet certain passenger-type mutations may pile up in greater numbers, if the underlying mutations are too easy to occur.

At nucleotide level, there are 12 routes of interchanges between A/G/T/C for single nucleotide substitutions, which constitute the vast majority of cancer mutations. The chances of generating each kind of mutations are certainly not equal. Many factors contribute to such phenomenon. First, different endogenous and exogenous mutagenic events lead to different types of nucleotide substitutions(Alberts B, Johnson A, Lewis J, 2002; Alexandrov et al., 2013, 2016; Farazi and DePinho, 2006; Kucab et al., 2019). Second, the abilities to recognize, repair and tolerate different types of mutations are also different(Frigola et al., 2017; Helleday et al., 2014). Third, although difficult to predict, different nucleotide sequences surrounding the mutation site may cause local variances that physically or chemically affect the ability for mutagens to attack. In addition, certain sequences are also more prone to be edited by enzymes such as APOBEC(Buisson et al., 2019; Nik-Zainal et al., 2012).Therefore, different flanking nucleotide sequences can also affect mutation rate(Alexandrov et al., 2013; Hodgkinson and Eyre-Walker, 2011; Ma et al., 2010).

Taken together, the probability to generate different types of nucleotide change may vary greatly. If two mutations both change the functional status of an important gene and promote cancer, they should be selected for during cancer formation. However, if one of such mutation is too difficult to generate at nucleotide level, the number of cancer cases carrying this mutation will significantly decrease. Considering this, if we can define the relative difficulty to generate each type of nucleotide mutations in cancer, we will be able to better estimate the functional importance of cancer mutations.

Although mutational signatures for ageing, UV, APOBEC, smoking and other cancer causes have been established, it is impossible to predict what percentage of cancers are influenced by each signature, and to what extent. Therefore, the relative difficulty to generate different types of mutation in cancer has not been established. In this report, through analysis of mutational data from 26,000 cancer genomes, we established the relative mutational difficulty for different types of cancer mutations, and showed that it can help accurately interpret functional importance of cancer mutations.

## Results

### Defining relative mutational difficulties in human cancer

Although it is very difficult to construct a mathematical model that could weight in all relevant factors to forwardly predict how much more difficult it is to generate one type of mutation versus the other, such differences do factually exist, and they collectively determined the mutation distributions in human cancer. Therefore, we argue that by analyzing large human cancer genome dataset, we can reversely derive the relative difficulties for each type of mutations (Fig. S1).

We retrieved mutation information for all human coding genes from the COSMIC database. From the approximately 26,000 cancer samples (Fig. S2) that were subjected to exome or whole-genome sequencing, more than 3 million single nucleotide mutations were identified on protein coding sequences (Fig. 1A). Considering that some mutations such as KRAS G12D and BRAF V600E are selectively enriched during cancer development, which could skew our estimation of mutation tendency, we excluded mutational events that occur in more than 5 cancer samples (see methods for discussion). This eliminated about 2% of mutations (Fig. 1A) and the remaining mutations were collated into different groups.

**Figure 1.**
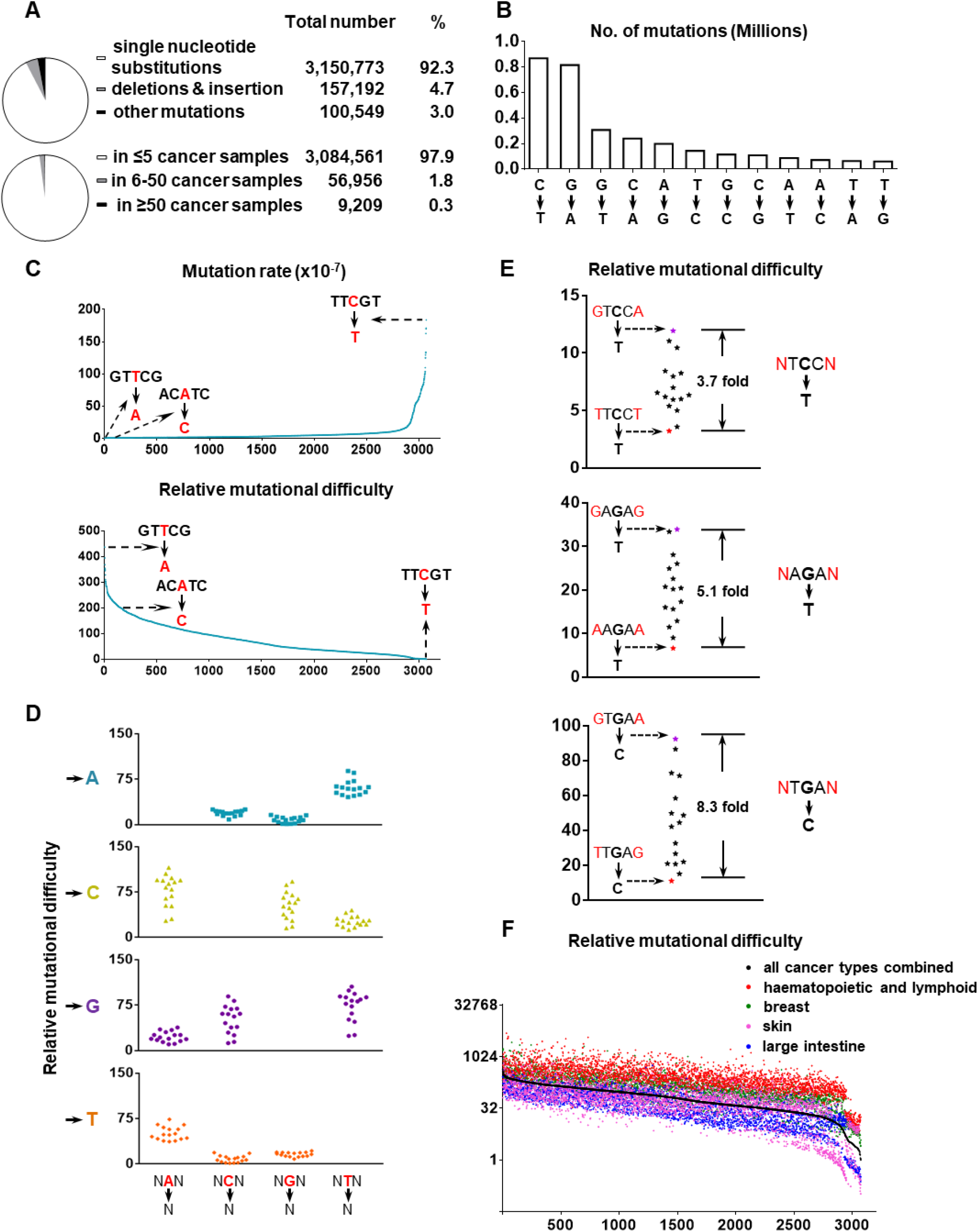
Relative mutational difficulty in human cancer. (A, B) Overview and classification of coding mutations from about 26,000 cancer genomes. (C) Rates and relative difficulties of different types of mutations based on 26,000 cancer genomes. Depending on the type of nucleotide substitution and the surrounding sequences, mutations are divided into 3,072 groups. (D, E) The impact of flanking nucleotides on relative mutational difficulty. (F) Cancer type-specific relative mutational difficulty.

Overall, the number of C→T mutations and its complementary G→A mutations constitute more than half of mutations in cancer (Fig. 1B). The rate of C→T mutation is 14 folds more than T-G mutation, demonstrating that the chances to generate each type of mutations do vary significantly (Fig. 1B).

Importantly, to reach a systematic view of how neighboring sequences might affect mutational tendency, we performed an extensive analysis, in which nucleotides at −2, −1, +1 and +2 position were all taken into consideration. Consequently, mutations were divided into 3072 groups (Table S1).

For example, the most likely to occur cancer mutation is C→T mutation on TTCGT sequences, which appeared 10,563 times. There are approximately 575 million TTCGT sequences in 26,000 coding genomes. Therefore, the chance of a C→T cancer mutation on TTCGT sequences can be calculated as 1.85*10^-5^ (=10,563/575,000,000), which is about 200 folds more than the probability of A→C mutation in an ACATC sequence (Fig. 1C). In other words, it is 200 times more “difficult” to generate the latter mutation in human cancer. Similarly, such “difficulty” indexes were generated for all 3072 types of nucleotide substitutions, which showed a wide distribution (Fig. 1C, S3 and Table S1). Analysis of these difficulty indexes showed that in addition to nucleotides on −1 and +1 positions (Fig. 1D), the nucleotides on +2 and −2 positions can also exert significant impacts on mutational tendency (Fig. 1E and Fig. S4). This indicates that it is important to incorporate the flanking nucleotide sequences into analysis when assessing individual mutations.

Our analysis shows that different types of mutations are generated at remarkably different rates (Fig. 1C). Given that the chance to generate different types of mutations can vary by several hundred folds, it strongly suggests the need to reassess human cancer mutations and our dataset will provide a useful tool.

To more precisely evaluate individual cancer mutations, we also took into consideration that certain types of human malignancies, such as melanoma, endometrial and colorectal cancers exhibit significantly higher mutation rates than other types of human cancer(Schneider et al., 2017). Therefore, the same type of mutation may be generated at significantly different rates in different types of cancer. Considering this, we generated cancer type-specific mutational difficulty indexes with similar method (Fig. 1F, Fig. S5 and Table S2), which will enable precise assessment of cancer mutations.

### Incorporating mutational difficulty to predict loss of function p53 mutations

We hypothesize that these “difficulty” indexes can serve as a valuable tool to predict the functional importance of cancer mutations. For example, if an A→C mutation in an ACATC sequence, despite the high difficulty, is still strongly selected for and appears in noticeable number of cancer samples, it could indicate that such a mutation is significantly enriched during cancer development. Therefore, such mutations may be crucial for cancer development. On the other hand, certain easy-to-occur passenger mutations may appear more common in cancer database, but their mutation counts may be more of a reflection of the high mutational tendency, rather than their importance.

We applied this method to assess the functional impact of p53 missense mutations. Several well-established p53 hotspot mutations account for about 27% of all p53 missense mutations and are known to abolish gene function. Most of the less frequent p53 missense mutations, although constituting the majority, are hard to predict in terms of their functional impact. We factored in the aforementioned “mutational difficulty” to forwardly estimate the functional importance of each mutation. For example, the M133R mutation is caused by T→G substitution on a GATGT sequence, whose difficulty index is 233. This mutation appeared in only 11 cancer samples in the COSMIC database. Given our argument, the frequency of this M133R mutation may have been severely penalized by the high mutational difficulty. Considering this, we designated M133R’s original count as 11 and revised count as 2563 (=11×233). Notably, the revised count for this mutation is comparable to that of the hotspot R282W mutation (original count 609, difficulty index 3.31, revised count 2017). Since such a revised mutation count integrates both the original count and mutational difficulty, it reflects the selective pressure for each mutation more closely. Consequently, it will provide a better reading of their functional impacts.

To more precisely assess these p53 mutations, we also took into consideration that the same type of mutations is generated at different rates in different cancer types (Fig. 1F). Therefore, in our analysis we used cancer type-specific mutational difficulty indexes to calculate the revised mutation count for each p53 mutation (see methods) (Table S3).

The global view of p53 missense mutations is provided in figure 2A. The map of p53 mutation original count is characterized by seven high peaks at R248 and R273, which are crucial for interaction with DNA, as well as R175, Y220, G245, R249 and R282, which are crucial for maintaining p53 structure. In the revised mutational count map, many more such high peaks appeared, suggesting that other portions of p53 also contain numerous amino acid residues that are essential for p53 function (Fig. 2A). Importantly, judging from original counts, only a few p53 missense mutations occur more frequently than the hotspot R282W mutation (Fig. 2B). After considering the mutational difficulty, more than 130 of p53 missense mutations exhibit a higher revised count than R282W (Fig. 2B), suggesting that many more p53 missense mutations potentially abolish gene function.

**Figure 2.**
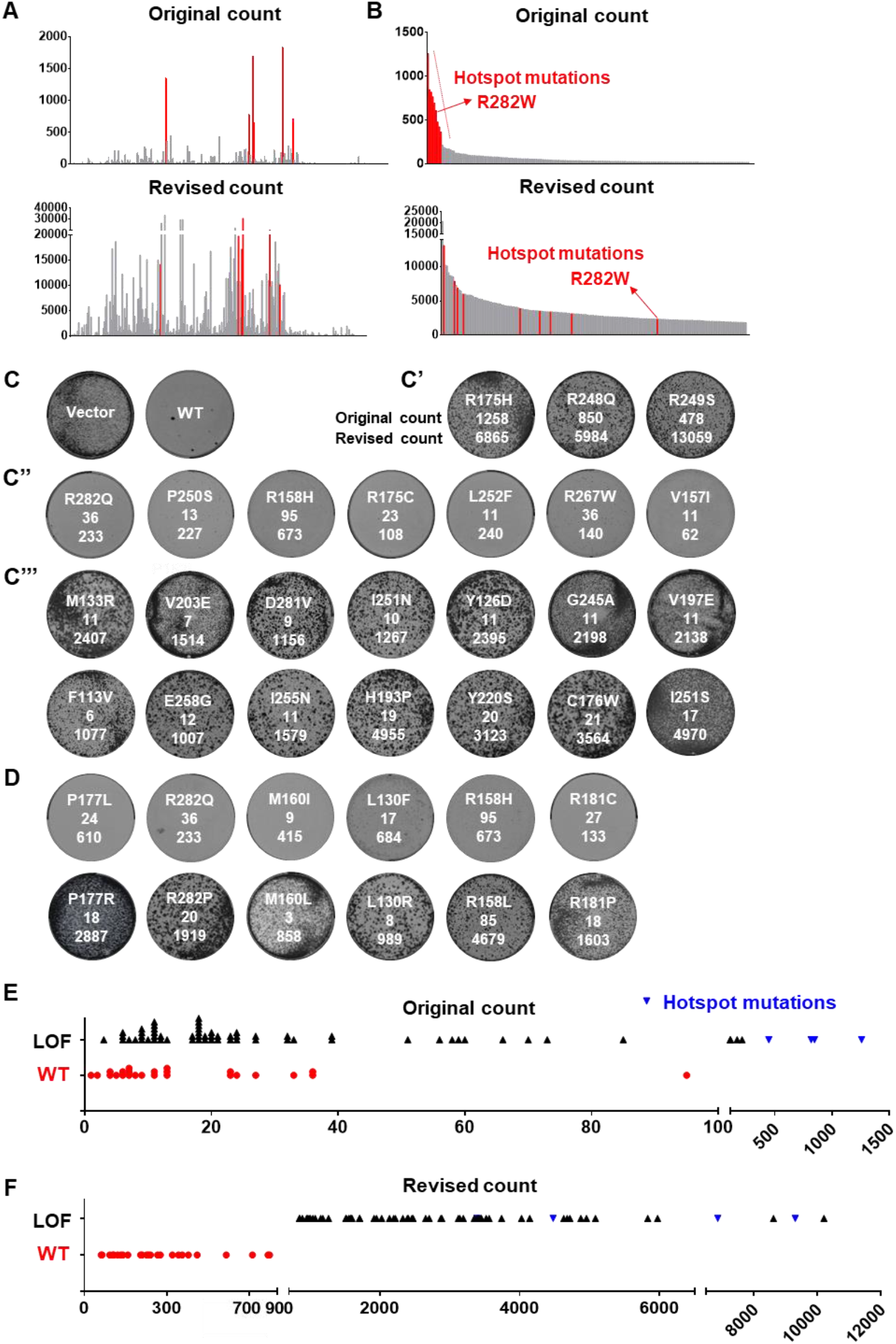
Integrating relative mutational difficulty to predict the functional status of p53 mutations. (A) p53 mutation histogram based on original and revised counts. Different types of mutations on the same amino acid residue (e.g. R273H and R273C) are combined to make the graph. Red lines indicate hotspot mutation sites such as G245 and R282. (B) The original and revised counts of p53 cancer mutations. Red lines indicate hotspot mutations such as R282W. (C) Expression of wild type p53 suppresses the growth of Saos-2 cells. Genes were delivered to cells via lentivral infection. For all colony formation assays in this study, cells were infected with low MOI such that 30-50% of cells infected with virus. (C’) p53 hotspot mutants are well-tolerated by Saos-2 cells. The original and revised counts are listed below each mutant. (C’’) p53 mutants with revised count lower than 700 behave like wild type p53 and suppresses Saos-2 growth. (C’’’) p53 mutants with revised count higher than 900 are loss of function mutants and are well-tolerated by Saos-2 cells. All colony formation assay in this study were done in three independent biological repeats. (D) Pairs of p53 mutations on the same amino acid. Shown are examples of high-difficulty mutations, although appearing in lower number of cancer samples, are loss of function mutations instead. (E) Original mutation counts do not correlate with functional status of p53 mutants. (F) Revised mutation counts correctly predict the functional status of p53 mutants.

To establish a cut-off value that could help identify p53 mutants that still retain wild type function, we compiled revised count values for all p53 synonymous mutations and found them to be mostly below 700 (Fig. S6). Therefore, a revised count below 700 may suggest wild type function for p53 mutants. We also estimated that a revised count over 900 might suggest loss of function. We constructed more than 80 low-frequency p53 mutants with various revised count values to test such hypothesis. The human osteosarcoma cell line Saos-2 carries homozygous deletion of p53. It could tolerate hotspot p53 mutants but not wild type p53 (Fig. 2C). Consistent with our hypothesis, all p53 mutants with revised count values of less than 700 behaved like wild type p53 in this assay (Fig. 2C), suggesting they do retain gene function as predicted by our method. For example, the R282Q mutation (original count 36) is located on the functionally essential amino acid residue R282. However, the underlying nucleotide substitution for this mutation is relatively easy to occur, and with a revised count of 186, this mutant retained wild type p53 function. We also noticed that the R158H mutation, although observed in 95 cancer samples, is an easy-to-occur mutation, and with a revised count lower than 700, this mutation also retained wild type function. The high number of cancer samples carrying this R158H mutation may be more of a result of the easiness of generating the underlying mutation.

In contrast, certain high difficulty p53 mutations, although many of which only occurring in less than 10 cancer samples, are predicted to be loss of function mutations with revised counts over 900. We examined around 50 of such p53 mutations, and they were all well tolerated by Saos-2 cells, confirming their loss-of-function status (Fig. 2C and Fig. S7). This is consistent with our hypothesis that for certain functionally important, but hard-to-occur mutations, their frequencies in cancer are greatly suppressed. Such mutations tend to be overlooked in previous studies. After factoring in mutational difficulty, we can predict the functional importance of such mutations.

We also noticed that, on P177, the P177L mutation (original count 24, revised count 262) retains wild type function (Fig. 2D). Interestingly, on the same residue is another mutation P177R, which exhibit lower original count but much higher revised count (original count=18, revised count=2887). Despite it being less frequent than P177L, it is actually a loss-of-function mutation (Fig. 2D). Importantly, in our analysis we observed multiple such cases that even on the same residue, less frequent mutations could be loss-of-function, yet mutants with higher original counts retain wild type function. Examples include R282Q/P, M160I/L, L130F/R, R158H/L and others (Fig. 2D). Such a reverse phenomenon can be explained by their revised mutation counts, again demonstrating the validity of our method.

To examine the biochemical function of these p53 mutants, we introduced them into HCT116 p53-/-cell line, and tested whether DNA damage drugs can still induce the expression of p21, a well-established p53 transcriptional target(el-Deiry et al., 1994). Again, p53 mutants with revised counts lower than 700 behaved similarly to wild p53, whereas p53 mutants with revised mutation counts higher than 900 behaved similarly to hotspot mutants, failing to upregulate p21 upon DNA damage (Fig. S8)

Summarized in figure 2E, despite the common perception that high-impact mutations appear more frequently in cancer database, the original mutation count is not a reliable predictor of functional status. p53 mutants with original counts less than 100 can either be loss of function mutants or retain wild type function. In contrast, the functional status of p53 mutants are accurately predicted by their revised mutation counts, with a value lower than 700 indicating wild type function, and a value higher than 900 indicating loss-of-function (Fig. 2F). This shows that by defining relative mutational difficulty, we can provide novel tools to accurately assess cancer mutations.

### Dominant negative effects of p53 mutants

It is known that human p53 hotspot mutations also exert dominant negative effect over wild type p53(Willis et al., 2004). To explore whether such dominant negative effect also exists for other p53 mutations, and whether our method could predict such dominant negative effect, we established an experimental system using the Eμ-Myc;p19Arf-/-mouse lymphoma cell line. This cell line retains wild type p53, which can be activated by DNA damage to induce cell death(Stott et al., 1998). Expression of hotspot p53 mutant together with GFP was achieved in this cell line via retroviral vectors. Hotspot p53 exerts dominant negative effects over endogenous wild type p53, and cells could not efficiently elicit cell death when treated with DNA damaging drugs. As a result, the percentage of GFP-positive, hotspot p53 mutant-expressing cells increased after drug treatment (Fig. 3A and 3B). In contrast, expression of wild type p53 in this system moderately sensitized cells to DNA damage drugs (Fig. 3B).

**Figure 3.**
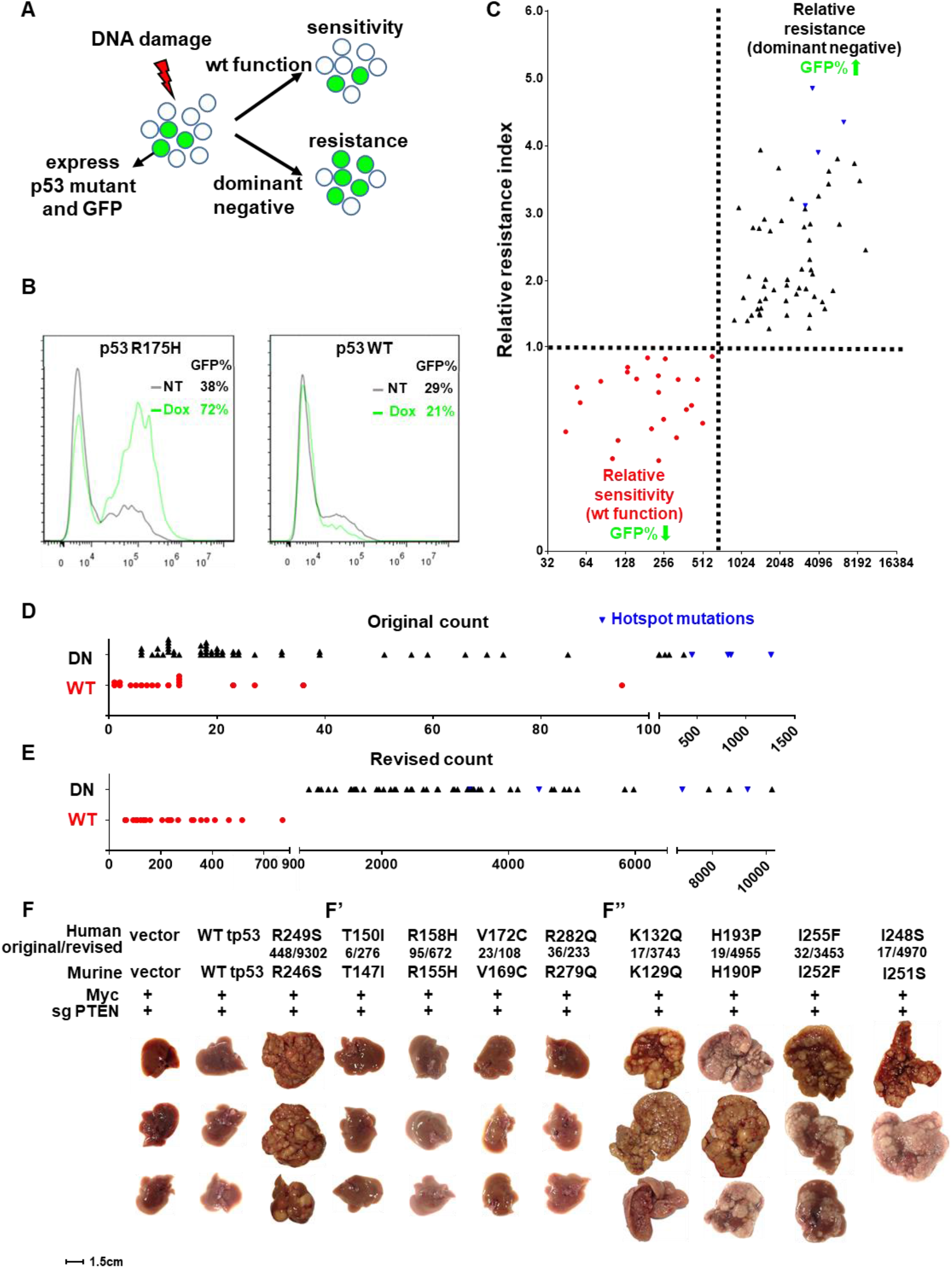
Integrating relative mutational difficulty to predict dominant negative effects of p53 mutations. (A) An experimental system to analyze dominant negative effects of p53 mutations. A murine lymphoma cell line that retain wild type p53 is partially infected by retrovirus that express p53 mutants and GFP. (B) If the p53 mutants exert dominant negative effect over endogenous wild type p53, it will render cells more resistant to DNA damage induced by doxorubicin, and the rate of GFP-positive cells increases in surviving cell population. Expression of wild type p53 moderately sensitizes cells to doxorubicin treatment. (C-E) Revised counts, but not original counts of p53 mutants correctly predicts whether such mutants exhibit dominant negative effects. Murine p53 mutants corresponding to human p53 mutants were used in these experiments. (F) Revised counts correctly predict whether p53 mutants can promote liver cancer formation in vivo. The original and revised counts are listed below each mutant, separated by a “/” mark. Murine p53 mutants corresponding to human p53 mutants were used in this experiment. Mice are sacrificed 30 days after hydrodynamic delivery of genes in vivo. n=3 for each experimental group, except I248S for which one of the injected mice didn’t recover from hydrodynamic injection.

We cloned the murine versions of various human p53 mutants and tested whether they exhibit dominant negative function. Importantly, all p53 mutants with revised counts lower than 700 behaved like wild type p53 (Fig. 3C), whereas all p53 mutants with revised count higher than 900 exhibited dominant negative effect (Fig. 3C). Again, as a predictor of dominant negative effect, revised count performed significantly better than original mutation count (Fig. 3, D and E).

We further tested whether our method can predict cancer-promoting abilities of p53 mutants *in vivo*. Using a tail-vein hydrodynamic injection method, together with a transposon system(Chen et al., 2007) and CRISPR gene editing (Yang et al., 2014), Myc overexpression and PTEN knockout was achieved in the liver of wild type FVB mice. Under such condition, no mice developed liver tumor at 3 weeks. Addition of R246S murine p53 mutant, which mimics the human R249S hotspot mutation, overrode endogenous wild type p53 in mice liver and caused massive tumors (Fig. 3F). Using this setting, we tested eight p53 mutants. Consistent with our prediction, four mutants with revised counts lower than 700, including the rather frequent R158H mutation (original count 95), all behaved like wild type p53 and caused no tumors. In contrast, four p53 mutants with high revised mutation counts all caused massive liver tumors in mice (Fig. 3F), indicating they were able to override endogenous wild type p53 to promote cancer.

### Functional importance of several regions on p53 structure highlighted by our method

Our results suggest there are many low frequency p53 mutations with significant functional impacts. To better understand their general distribution, using the revised counts of p53 hotspot mutation sites as controls, we mapped these high-impact mutations on the three-dimensional structure of p53. Known p53 hotspot mutations are located on the interfaces that are crucial for p53 structure and interaction with DNA. We first noticed that many residues adjacent to hotspot sites, such as V173, H178, M246, V274 and A276, although with rather low original mutation counts, showed very high revised counts (Fig. 4A and Fig. S9A). This observation suggests that many residues surrounding hotspot sites are in fact also essential for p53 function. These crucial sites are rarely mutated in cancer, because their mutation frequencies are severely penalized by high mutational difficulty.

**Figure 4.**
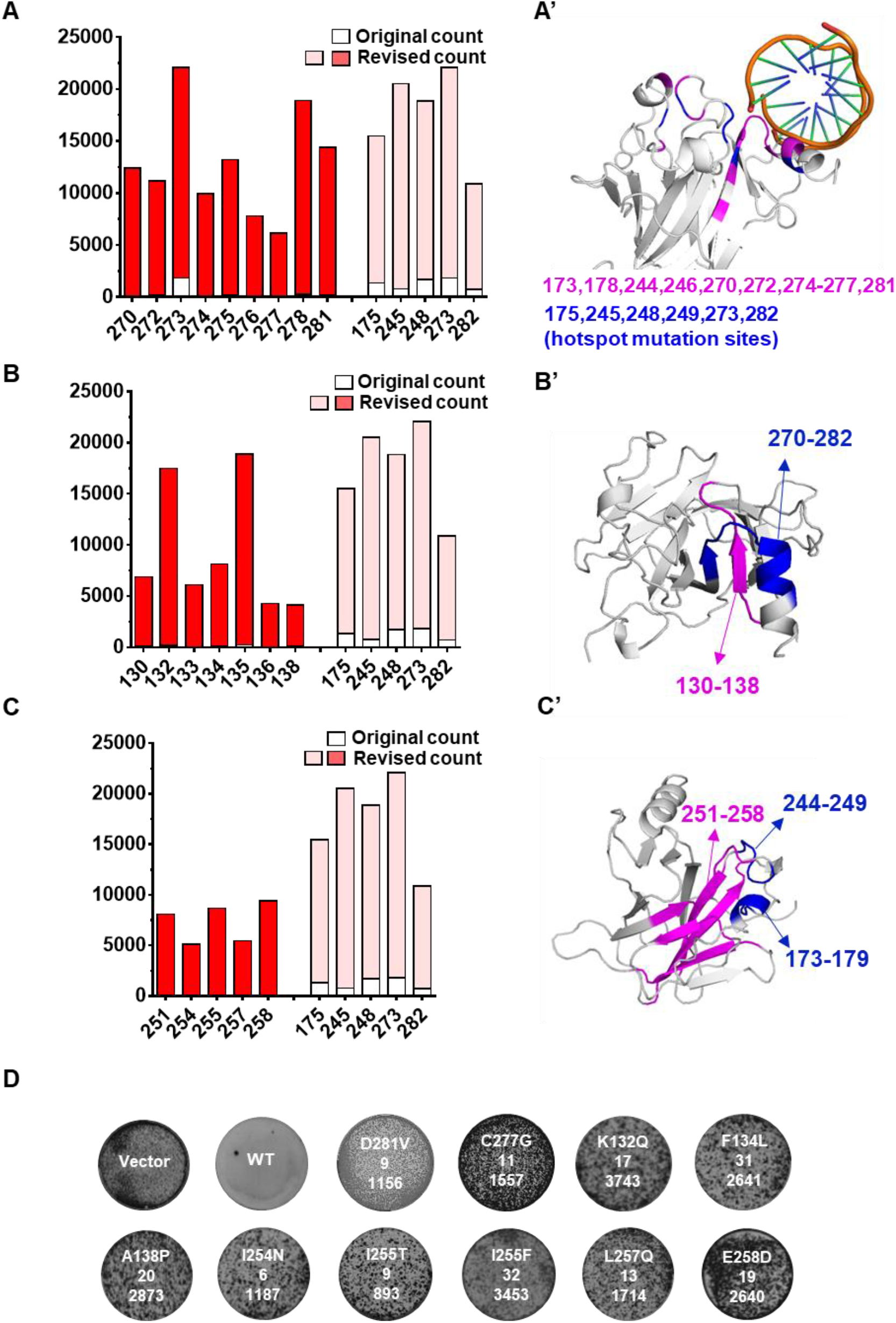
Functionally important amino acid residues and regions in p53. (A-C) On the left panels, original mutation counts of listed amino acid residues are indicated by white boxes, whereas revised counts are indicated by red or pink boxes. Hotspot mutation sites such as R175 and G245 are included as controls. On the right panels, (A’) Functionally important amino acid residues near hotspot mutation sites are labeled in purple. Hotspot mutation sites are labeled in blue. (B’) and (C’) Additional regions crucial for p53 function are labeled in purple. Regions that mediate DNA binding such as AA270-282 and AA173-179 are labeled in blue. (D) Example of low frequency mutations on amino residues in regions analyzed in (A-C). These mutations abolish p53 function and are well tolerated by Saos-2 cells.

In addition to these residues, several other regions of p53 stood out with high revised mutation counts. One of such rarely-mutated, high impact region is residues 130 to 138, which form a β strand and loop structure that lays closely to a β strand-loop-helix domain (amino acids 270-282), which host several hotspot mutations and are responsible for DNA-interaction(Eldar et al., 2013; Follis et al., 2014) (Fig. 4B). Other high impact amino acids identified by this method are five β strands that formed the central β-barrel of p53. On the three-dimensional structure of p53, such amino acids are also very close to AA173-179 and AA244-249, both hosting hotspot mutations(Walker et al., 1999) (Fig. 4C and Fig S9B). Colony formation assays in Saos-2 cells confirmed that many rare, but difficult-to-generate mutations on these sites disrupt p53 function (Fig 4D). Such mutations were also tested in the Eμ-Myc;p19Arf-/-system and exhibited dominant negative effects over wild type p53 (Fig 3C). Taken together, our method is able to regroup p53 mutations by integrating mutational difficulty, and points to additional regions that are crucial for the function of p53.

### Predicting the functional status of PTEN and INK4A mutations

Next, we asked whether this method could be applied to other established cancer genes such as PTEN and INK4a. We cloned about 20 low-frequency PTEN and INK4A mutations and expressed them in PTEN or INK4A deficient cancer cell lines to see whether such mutations abolish gene function. As predicted by our method, those mutations with low revised counts retained wild type function, whereas those mutations with high revised counts caused gene loss of function (Fig. 5A-C).

**Figure 5.**
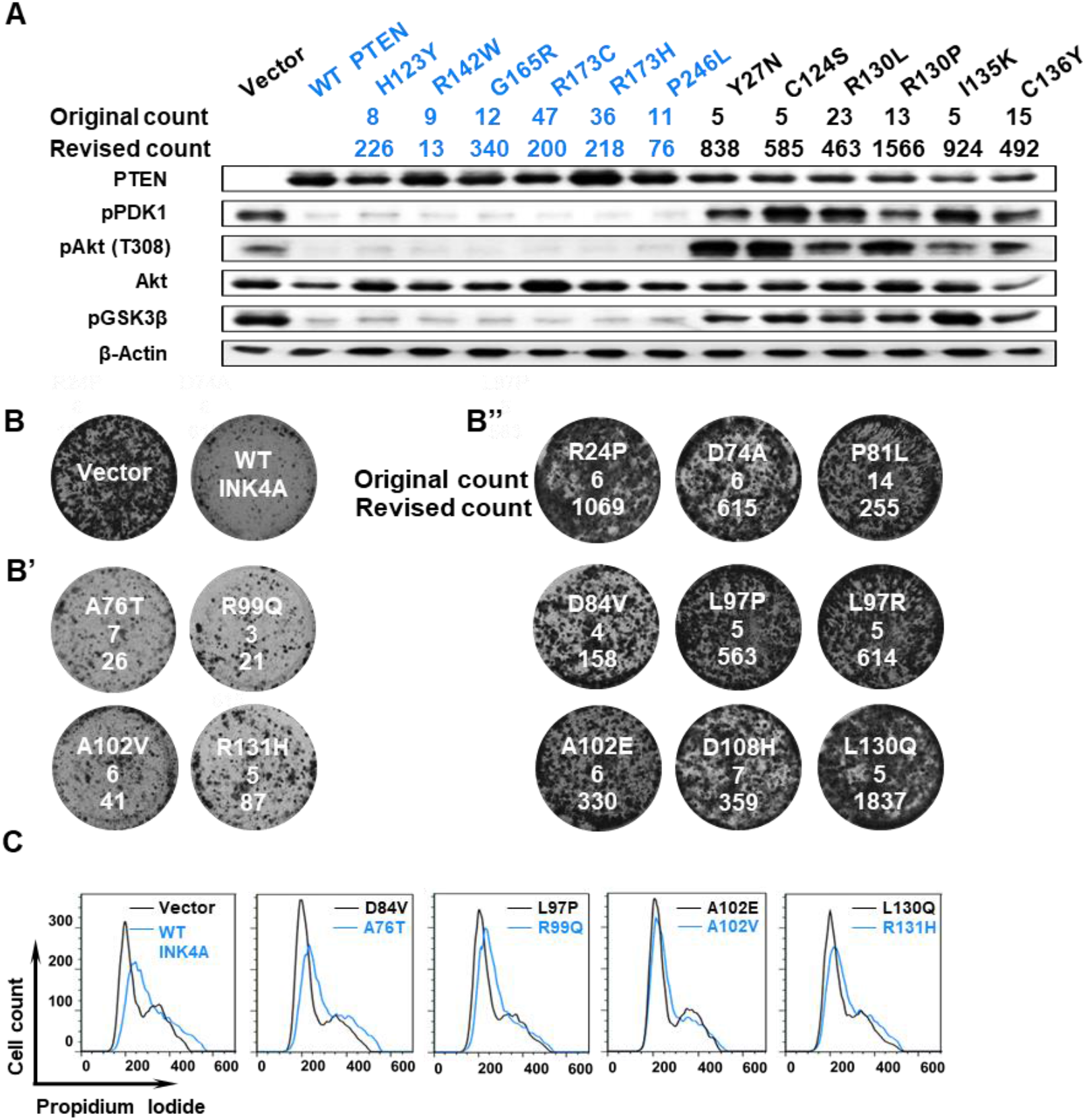
Integrating relative mutational difficulty to assess the functional status of PTEN and INK4A mutations. (A) Original and revised counts of indicated PTEN mutants. Expression of wild type PTEN suppresses AKT signaling in in 786-O cells, which do not express PTEN. PTEN Mutants with low revised counts (in blue) behave like wild type PTEN and are able to suppress downstream AKT signaling. PTEN mutants with high revised counts (in black) lose PTEN function and cannot suppress AKT signaling. The original and revised counts are listed below each mutant. (B) Expression wild-type INK4A suppress growth of U251 cells, in which the INK4A locus is deleted. B’) INK4A mutants with low revised counts retain wild type gene function and suppressed U251 growth. B’’) CDKN2A mutants with high revised counts are defective in gene function and are well tolerated by U251 cells. The original and revised counts are listed below each mutant. (C) INK4A mutants with high revised counts (in black) cannot induce cell cycle arrest in U251 cells. CDKN2A mutants with low revised counts (in blue) behave like wild type CDKN2A and induce G1/early S phase arrest in U251 cells.

For example, high difficulty mutations including PTEN Y27N, C124S and I135K, although each only occurring in 5 cancer samples in COSMIC, could not suppress AKT signaling, proving that they all abolish PTEN function (Fig. 5A). In contrast, the PTEN R173C and R173H mutations, despite being the fifth and sixth most common PTEN mutations and occurring in 47 and 36 cancer samples, both retained wild-type function (Fig. 5A). According to our method, both are low-difficulty mutations, which explains why they did not disrupt PTEN function despite being detected in significantly more cancer samples. This observation, together with finding that p53 R158H mutation (original count 95, revised count 673) also retains wild type function, demonstrate that our method not only help identify rare mutations that promotes cancer, it can also point out high frequency, passenger type mutations in cancer database.

### Potential implications for cancer prevention

In addition to establishing a method that could help evaluate individual cancer mutations in a sequence- and cancer type-specific manner, we also asked whether our method can help understand cancer etiology in general. In our analysis of human cancer mutations, we noticed that the vast majority of common mutations on tumor suppressors, such as p53, PTEN, FBXW7 and SMAD4 are low-difficulty mutations (Fig. 6A). 77% of these common, low difficulty mutations are C to T or its complementary G to A mutations on CpG sequence, which could be the results of spontaneous deamination(Rideout et al., 1990). From a cancer prevention point of view, it will be rather hard to prevent such type of deleterious events on tumor suppressors.

**Figure 6.**
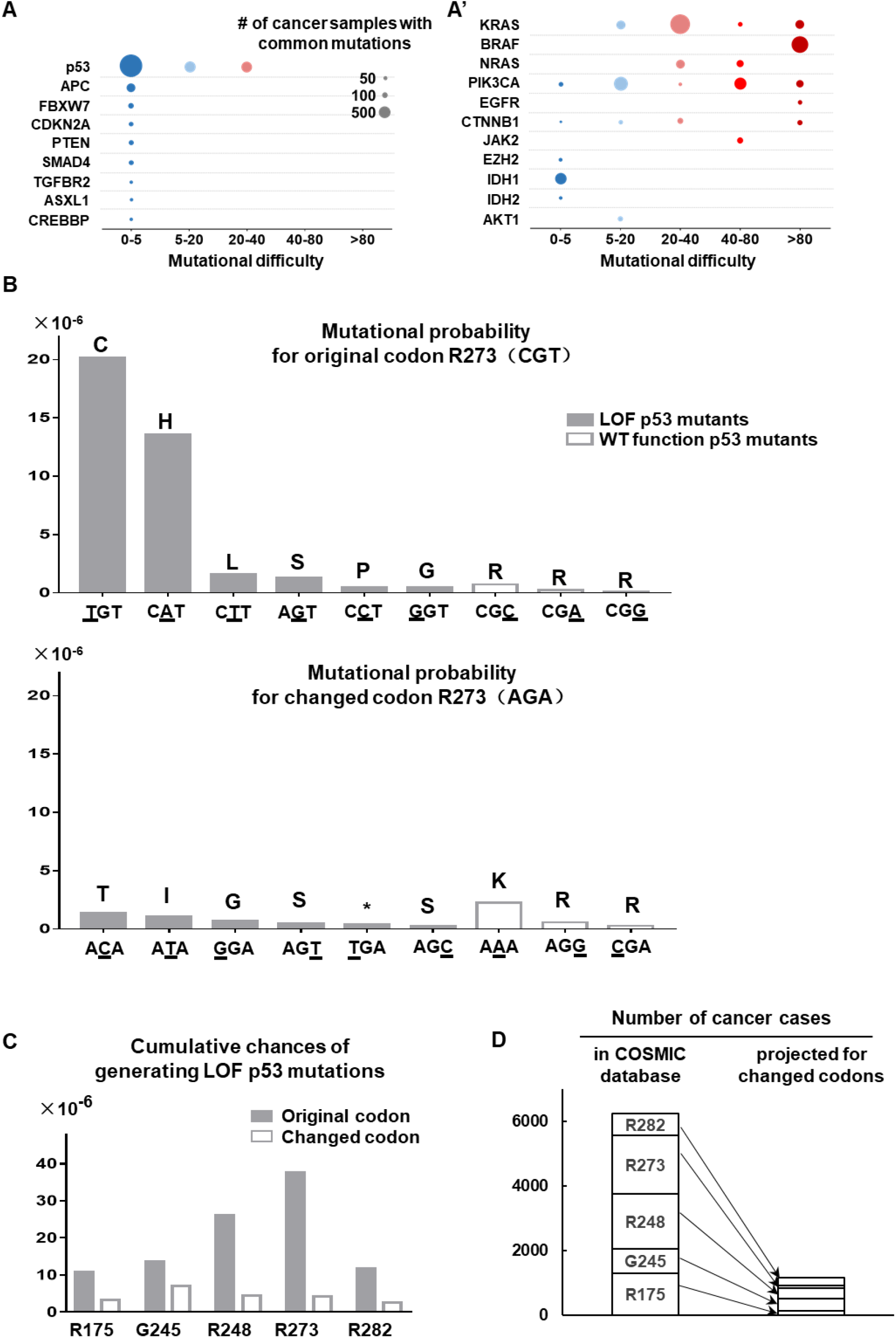
Potential implications for cancer prevention. (A) and (A’) Relative difficulties of common mutations on established oncogenes and tumor suppressors. Mutations on genes such as KRAS and p53 are collected from the about 26,000 cancer genomes. Those mutations appearing in more than 30 cancer samples are plotted according to their relative mutational difficulties. For example, in (A’) the blue dot adjacent to p53 represents how many cancer samples carry common p53 mutations whose relative mutational difficulties are between 0 and 5. (B) Mutational probability of the p53 R248 original codon R(CGG) and changed codon R(AGA). Loss of function mutations are shown in dark grey. Mutations with wild type p53 function are shown in white. (C) Cumulative chances of generating LOF p53 mutations on five hotspot mutation sites, based on original and changed codons. (D) Number of cancer cases in COSMIC database involving these five p53 hotspot sites, and the projected reduction of cancer cases if these sites are changed to hard-to-mutate codons.

In contrast, except for IDH1/2 and AKT1, common mutations on oncogenes such as KRAS, BRAF, CTNNB1, PI3KCA and JAK2 are typically high-difficulty mutations (Fig. 6A’). Only 14% of common cancer-promoting mutations on oncogenes are C to T or G to A mutations on CpG sequences. About half of common cancer-promoting mutations on oncogenes require purine to pyrimidine changes or vice versa. Such type of changes cannot be caused by simple chemical reactions such as deamination of the nucleobase. Rather, Strong exogenous carcinogenic events are probably needed to damage DNA to eventually create such types of mutation. Limiting the exposure to environmental carcinogens, as well as managing long-term inflammation, among many potential measures, may significantly reduce the chance of obtaining activating mutations on oncogenes in general. This will significant deplete the driving force of cancer and impede cancer development. Therefore, even though two third of mutations in human cancer are caused by spontaneous events (Tomasetti et al., 2017), avoiding environmental carcinogenic factors holds great promises to significantly reduce the incidence of many types of cancer.

On the other hand, certain types of cancers will still be hard to prevent. For example, the driver mutations on the oncogenes IDH1 and IDH2 are both low-difficulty, C to T or G to A mutations (Fig. 6A’). Therefore, cancer cases associated with such mutations, for example subtypes of glioblastoma, cholangiocarcinoma and acute myeloid leukemia (Borger et al., 2011; Dang et al., 2010) may be hard to prevent. Similar to the argument by Tomasetti et al (Tomasetti et al., 2017), for these types of cancer early detection holds more promise than cancer prevention methods.

Lastly, it is apparent that low difficulty, spontaneous mutations on tumor suppressors contribute to human cancer, most significantly through several easy-to-mutate sites on p53 (Fig. 6A). From a pure theoretical point of view, it is possible to introduce synonymous mutations to these sites to render them more resistant to deleterious mutations. We analyzed the potential benefits of changing the nucleotide coding sequence on p53 hotspot sites. For example, changing the p53 R273 sequence from R(CGT) to R(AGA) will reduce the chance of generating loss of function mutations on this site by seven folds (Fig. 6B). For four other mutational hotspots on p53, similar codon changes can also significantly reduce the chance of generating loss of function mutations (Fig. S11) and are projected to greatly reduce cancer cases involving these hotspot sites (Fig. 6C-D). If theoretically, spontaneous low-difficulty mutations on p53 can be limited by such measures, and high-difficulty mutations on oncogenes and other sites of p53 can be thwarted by avoiding environmental carcinogens, it may dramatically reduce cancer incidence.

## Discussion

### Significant differences of mutational difficulties in human cancer

The functional importance of a mutation to cancer can be reflected by its selective enrichment in cancer samples. However due to the lack of understanding of relative mutational difficulty in cancer, most studies use mutation frequency in cancer database to directly calculate selective pressure. Our analysis shows that, depending on the type of nucleotide substitution and the surrounding sequences, the chances of generating different types of mutations can vary by as much as 400 folds (Fig 1C). Such a drastic difference highlights the need to re-approach how we calculate the selective pressure for each cancer mutation, and consequently how to interpret the functional importance of such mutations.

Many factors contributed to the fact that different types of nucleotide substitutions are created at rather different rates(Alberts B, Johnson A, Lewis J, 2002). Given the complex nature of these processes, relative mutational difficulties for different types of mutations, as well as their impact on how to interpret human cancer genome have not been established. In this report, through analysis of large number of human cancer genomes, we reversely derived the relative difficulties for each type of mutation. We also established such numbers in a cancer type-specific manner. Such dataset will be a useful tool to understand cancer genome.

For most genes, close to 30,000 cancer samples have been analyzed and deposited in the COSMIC database. Certain easy-to-occur mutations may simply accumulate in a number of cancer samples without provide advantages for cancer development. In the future, when increased number of cancer genomes are deposited to COSMIC database, it is expected that more and more easy-to-occur passenger mutations will pile up on the mutation histogram. Without considering relative mutational difficulty, these seemingly “mutational peaks” may lead to erroneous assumptions that they are cancer-driving mutations.

To functionally estimate the importance of novel cancer mutations, the cancer types that host such mutations should also be taken into consideration. As illustrated in figure S2, for many types of mutations, it is much easier to generate them in skin, colorectal and endometrial cancers. On the other hand, some types of mutations are more difficult to generate in these cancer types (Fig. S3A). Therefore, if the original count of a cancer mutation is primarily contributed by skin, colorectal and endometrial cancer samples, such mutations should be viewed with caution. However, they should not be automatically devalued either.

### Functional landscape of p53 mutations in human cancer

Our results suggest that, in addition to hotspot mutation sites, there are numerous other amino acid residues also crucial for p53 function. Mutations on these residues are rare because it is too difficult to generate inactivating mutations on them. By incorporating relative mutational difficulty, we can re-establish the functional importance of such sites and discover many rare, but cancer-promoting p53 mutations. We also observed dominant negative effects for many p53 missense mutations (Fig. 3C), which is consistent with recent findings by Boettcher et al(Boettcher et al., 2019). This led to recognitions of amino acid residues and structural regions crucial for p53 function (Fig. 4), which will help further understand this important tumor suppressor.

Based on our analysis, we estimate that out of the 1219 types of missense p53 mutations in COMIC database, 27% are loss of function mutations and 70% retain wild type function. In addition, out of the 19598 p53-mutated cancer samples in COSMIC database, 83% contain loss of function p53 mutation and 15% retain wild type p53 function.

The FATHMM(Functional Analysis through Hidden Markov Models) method(Shihab et al., 2013), which estimate functional impact based on sequence conservation and the overall tolerance of the protein/domain to mutations, has been commonly used to predict cancer-driving mutations. Such a method is used by COSMIC to annotate cancer mutations. With regards to p53 mutations, comparison of our prediction results with FATHMM method showed only 50% overlap (Fig. S10 and Table S4).

Interestingly, in a recent publication, thousands of different types of p53 mutants were introduced to cancer cells, and the functional status of these p53 mutations were assessed by whether these mutations were tolerated by cells under different conditions(Giacomelli et al., 2018). Such dataset provides direct experimental readout of p53 mutations. Our predictions of the functionality of p53 mutations are highly similar to these experimental results, with a 88% overlap (Fig. S10). The 12% p53 mutations that are differently predicted (Table S5) may have come from wrong predictions by our method, or from small inaccuracies of such pool-based large-scale studies.

In the Giacomelli et al report, integrating mutational signatures of aging, APOBEC, smoking, UV, mismatch repair deficiencies and alatoxin, a model was trained to fit the mutational landscape of p53 in human cancer. In that model, for each p53 mutation, different sets of weights were assigned for each signature. Our analysis, by defining relative mutational difficulties, provides an straightforward method to predict the funtional importance of p53 mutations. This method can also be readily applied to PTEN, INK4A (Fig.5) as well.

In summary, our study provided estimates of relative mutational difficulties in human cancer. Such information may help further understand cancer formation, bringing new tools for discovering cancer-promoting mutations. Moreover, such kind of analysis may also yield additional perspectives on cancer prevention.

## Methods and Material

### Data acquisition

Mutation data from 26,154 cancer genomes were retrieved from COSMIC website in January 2018. If a gene has multiple isoforms, only the major form was included in our analysis such that mutations on the same sites are not counted multiple times. For 19,940 genes, 3,101,161 single nucleotide substitutions were identified. In order to calculate the natural mutational tendency, we first eliminated mutational events that occur more than 5 times in the dataset. These mutations may have been selectively enriched during cancer development and may interfere with our calculation. A total of 65,594 mutational events involving 6,019 sites were removed, which represent 2% of mutations. Detailed analysis of this point is provided in figure S2.

To assess the influence of neighboring sequences on mutational tendency, sequences of the coding genome corresponding to the 19,940 genes were downloaded from Ensemble (GRCh38.p10). For each nucleotide mutation, −2, −1, +1, +2 nucleotides were extracted from the corresponding coding sequence.

### Types of mutations

At the central position, there are 12 routes of interchange between A/G/T/C. The permutations at −2, −1, +1, +2 nucleotides amount to 4^4^. Therefore, we collated all mutations into 12×4^4^=3072 groups.

### Calculation of Mutational tendency

For the aforementioned 3072 groups, we first counted how many mutations from the 26,154 cancer genomes belong to each group. Next, we counted how many times each penta-nucleotide sequence appears in coding sequences of the 19,940 genes. For example, there are 10,389 C→T mutations on TTCGT sequences in 26,154 coding genomes. There are 21446TTCGT sequences per coding genome. Therefore, the mutational tendency of C→T on TTCGT is approximately 10389/ (21446×26154) =1.85 ×10^-5^, which is the highest amongst all 3072 combinations. We set the “difficulty” score for such a mutation as 1. The mutational tendency of A→C mutation in a CGATG sequence is 0.93×10^-7^, and its relative difficulty score is calculated as 1.85×10^-5^/0.93×10^-7^ = 200. Difficulty scores for all other combinations were generated accordingly.

In the above analysis, we aim to estimate the natural mutational tendency for each type of mutation in human cancer. Certain cancer-promoting mutations on genes such as KRAS and BRAF are strongly selected for during cancer formation. The number of such mutations are significantly increased in the dataset, not because they are easy to generate, but because they are strongly enriched by the tumorigenesis process. Therefore, their presence in the dataset may skew our estimation of the natural mutational tendency for each type of mutation. Considering this, in the above calculation, we excluded mutations that occur in more than 5 cancer sample, in order to achieve a closer estimate of mutational tendency. Of note, about 2% all of mutations in the 26,000 cancer genomes (Fig. 1A) occur in more than 5 cancer samples and were excluded in our analysis.

Fig.S3 shows the comparison of mutational difficulties calculated with and without excluding such recurrent mutations. In Fig.S3A, if no mutations are excluded, the mutational difficulty scores for KRAS G12R, BRAF V600E and HIF1A K213Q, among many others, will be significantly lower. In Fig. S3B, if only excluding mutations that occur in more than 20 samples, the mutational difficulty scores for TP53 V157G, NOTCH1 D573A, CDKN2A A36G, among others, will still be significantly lower. In Fig. S3C, if only excluding mutations that occur in more than 10 samples, the mutational difficulty scores for TP53 Y126D, KDM6A T794P, PIK3CA V344G, among others, will still be noticeably lower. Based on this, we calculated mutational difficulty after excluding mutations that occur in more than 5 samples.

Lastly, the numbers of different penta-nucleotides in the coding genome vary greatly. For example, in the coding genome there are 3,001 TAGCG sequences and more than 100,000 TGGAG sequences. Therefore, it is necessary to divide the number of mutations by the number of available sites to accurately understand the relative mutational difficulty.

### Cancer type-specific mutational difficulty

To generate cancer type-specific mutational difficulty scores, mutations were first grouped by cancer types, from which mutation rates were calculated using similar methods. For example, we observed 1,195 C→T mutations on TTCGT sequences in 296 endometrial cancer samples, and the mutational tendency of C→T on TTCGT is approximately 1.88×10^-4^ in endometrial cancer. Since in previous calculation we set the mutational difficulty score as 1 for a mutation rate of 1.85 ×10^-5^, we can calculate the relative mutational difficulty for C→T on TTCGT as 0.1 (=1.85 ×10^-5^/1.88×10^-4^) in endometrial cancer. This method was also used to calculate mutational difficulty for other cancer types.

### Analysis of p53, PTEN, INK4A mutations

Mutational data for p53, PTEN and INK4A was last acquired from COSMIC on January 2018. At the time, p53 was sequenced in 130,448 cancer samples, PTEN in 72,199 samples and INK4A in 72,566 samples. Current numbers in COSMIC database have slightly increased due to website updates.

Of note, the CDKN2A locus contains two genes, INK4A and ARF. Previous studies showed that recurrent mutations on the CDKN2A locus do not change the function of the ARF gene(Quelle et al., 1997). In addition, U251 cells, which deleted the CDKN2A locus, could tolerate ARF expression, but not INK4A expression. Therefore, for later experimental validation, we cloned and analyzed INK4A mutants in this study.

For each mutation, we first extracted the penta-nucleotide sequence surrounding the mutation site, and matched it with relative mutational difficulty scores. For mutational sites that are adjacent to intro-exon junctions, the genomic sequence was used to extract the nucleotide sequences surrounding the mutational site.

Next, we calculated the revised mutational count based on original mutation count and cancer type-specific relative mutational difficulty. For example, if a p53 mutation occurs in 10 colorectal cancers and 5 lung cancers, and the relative mutational difficulties for the mutation is 1 and 3 in colorectal and lung cancer respectively, the revised count for such a mutation can be calculated as 10×1 + 5×3 = 25.

Figure S6 shows the distribution of revised mutation counts for all p53 synonymous mutation based on COSMIC data. For most of these synonymous mutations, the revised counts are below 700. Therefore, we estimate that those missense p53 mutations with revised count below 700 retain wild type p53 function, which were validated with further experiments. In Figure S6, we also observed that the revised counts of several p53 synonymous mutations exceeded 700. This is due to the fact that certain p53 synonymous mutations abolish p53 function. For example, the p53 T125T mutation (c.375 G to A/C/T) disrupts the adjacent intron-exon splice site, and abolishes gene function(Supek et al., 2014). The revised count for T125T is 1353, and is predicted to be a loss of function mutation by our method.

Table S3 listed the original and revised counts of p53 mutations based on COSMIC database. If a mutation’s revised count is lower than 700, it is predicted to retain wild type function. If a mutation’s revised count is higher than 900, it is predicted to be loss of function mutation. A few exceptions exist and are explained below.

In the COSMIC database, the S149F mutation on p53 is caused by single nucleotide substitution in 5 samples, and the revised count is lower than 700. However, in one additional cancer sample, a CC to TT nucleotide change also caused the S149F mutation. Because of the rarity of such double mutations, we did not assign relative mutational difficulty score to such double mutations. Therefore, we cannot make functional prediction for this mutant, and a “*” is marked in the “revised count” column for S127F. Such phenomenon also occurred for S166L, V218M and R158C, and these mutations are labeled similarly with a “*” in Table S3. Several other mutations (for example R282W) also exhibited such double nucleotide substitution, however their revised counts calculated from single nucleotide substitutions already exceeded 900. Therefore, such mutations are predicted to be loss of function mutations in Table S3.

### GFP-based cell survival competition assay to determine sensitivity change caused by p53 mutants

The experiment was carried out with a protocol modified from(Bruno et al., 2017). Briefly, Eμ-Myc p19Arf−/− cells are infected with retrovirus that express GFP and mutant p53, such that 20-50% of cells are GFP positive. Cells are treated with DNA damage drug at doses that would kill 80-90% of uninfected Eμ-Myc p19Arf−/− cells. In this assay, if p53 mutant exerts dominant negative effects on endogenous wild type p53 in Eμ-Myc p19Arf−/− cells, after DNA damage drug treatment the GFP positive, p53 mutant-expressing cells will be relatively more resistant than GFP negative cells that only express wild type p53. At 72 h, treated and untreated cells are analyzed by flow cytometry. GFP percentages of live (PI-negative) cells are recorded and used to calculate relative resistance index.

### Calculation of relative resistance/sensitivity from GFP-based cell survival competition assay

The value of relative resistance index (RI) can be calculated as RI=(G2-G1*G2)/(G1-G1*G2). G1 means how many percentages of cells are GFP positive before drug treatment. G2 means how many percentages of cells are GFP positive after drug treatment. The explanation for such calculation was provided in (Jiang et al., 2011).

Relative resistance index larger than 1 means the corresponding p53 mutant displayed dominant negative effect, protected cells from DNA damage, and the rate of GFP+ cells in surviving cells increased after drug treatment. Relative resistance index smaller than 1 means the corresponding p53 mutant displayed wild type function, sensitized cells to DNA damage, and the rate of GFP+ cells in surviving cells decreased.

### Cell lines and drugs

Eμ-Myc p19Arf −/− cell was cultured in B-cell medium (45% Dulbecco’s modified Eagle’s medium and 45% Iscove’s modified Dulbecco’s media, supplemented with 10% fetal bovine serum, L-glutamate, and 5 μM β-mercaptoenthanol). Phoenix, HCT116 p53-/-, Saos-2, U251, A549, 293T and 293A were cultured in Dulbecco’s modified Eagle’s medium supplemented with glutamate and 10% (v/v) FBS. 786-O cell was cultured in RPMI medium supplemented with glutamate and 10% (v/v) FBS.

Saos-2, HCT116 p53-/-, U251, 786-O cells were obtained from the Cell Bank, China Academy of Sciences (Shanghai, China). Doxorubicin was purchased from Selleck.

### Antibodies

Antibodies against Phospho-Akt (Thr308) (D25E6) (Cell signaling, #13038), Akt (pan) (C67E7) (Cell signaling, #4691), Phospho-GSK-3β (Ser9) (D84E12) (Cell signaling, #5558), Phospho-PDK1 (Ser241) (C49H2) (Cell signaling, #3438), PTEN (pan) (Y184) (Abcam, #32199) were used for Western blot analysis.

### Cloning of p53, PTEN and INK4A mutants

Wild-type p53, INK4a and PTEN expression vectors were constructed as follows. The full-length open reading frame of p53, INK4a and PTEN cDNAs were amplified by PCR using KOD plus neo DNA polymerase (Code No.KOD-401 Lot No.646300) and a pair of primers with EcoRI and XhoI sites. The PCR product was cloned into the EcoRI/XhoI sites of the pMSCV-IRES-GFP vector. cDNAs with missense mutations were constructed by overlap extension PCR. All of the plasmids were sequenced to confirm that the appropriate mutations had been incorporated and that no additional mutations were generated.

All p53, INK4a and PTEN mutants tested in this study are listed in Table S6.

### Colony formation assay

To test the functional status of p53 mutants, retrovirus that expresses p53 mutants, puromycin resistance gene and GFP was used to infect Saos-2 cells. Cells are infected with similar virus MOI such that 30-50% of cells are GFP positive for all experimental groups. 5000 GFP-positive Saos-2 cells were resuspended in medium containing 10% FBS and plated in 6-well plates. After 24 hours, they were treated with 2ug/ml puromycin. 24 hours later, puromycin-containing medium was replaced with fresh complete culture medium. 5 days later, 2ug/ml puromycin was again used to treat cells for 24h before removal. Cells are cultured for an additional 10 days. Colonies were then fixed with 4% paraformaldehyde and stained with 0.1% crystal violet for 30 min. Stained cell colonies were washed with phosphate buffered saline (PBS) for three times and dried. Images were obtained by a digital camera. Similar protocols were used to test INK4a mutants in U251 cells.

### Mouse liver cancer model

Myc cDNA and p53 mutants were cloned into a transposon system using the PT3 vector(Chen et al., 2007). Such plasmids were mixed with sgPTEN-Cas9 plasmid(Yang et al., 2014), together with Sleeping Beauty transposase-expressing plasmid in PBS. Gene mixture was delivered to mouse by tail vain hydrodynamic injection. A rapid tail vein injection protocol, which delivers 2ml of PBS-plasmid solution in 7 seconds, introduced these plasmids into mouse liver cells. Under such conditions, PTEN is disrupted by sgPTEN and Cas9, and cDNAs of Myc and p53 mutants are integrated into host cell’s chromosomes by the transposon system. Concentrations of Sleeping Beauty transposase and Myc-expressing plasmids were at 0.5μg/ml and 1.25μg/ml, respectively. Other plasmids or corresponding empty vectors were used at 5μg/ml. Wild type FVB mice were used in this experiment.

### Cell cycle analysis

U251 cells expressing wild type or mutant forms of INK4a were analyzed. When grew to proper density (about 70-80%), cells were collected and fixed overnight in 70% ethanol. Cells were then treated with 0.2% Triton X-100, 50 μg/ml propidium iodide and 100 μg/ml RNase A for 40 minutes, and analyzed by FACS.

### Quantitative real-time PCR assay(qPCR)

RNA was purified using GeneJET RNA Purification Kit (thermo scientific) and qPCR was performed on a StepOne real-time PCR machine (BIO-RAD) using SYBR Green PCR master mix (Promega). mRNA level of actin was used as control. Primers used for qPCR analysis are listed in Table S8.

### Statistics

Differences of event frequency between two groups were analyzed using Student’s unpaired two-tailed T test. p-values <0.01 were marked as *** in figures, p-values <0.05 were marked as ** in figures.

## Author Contributions

Hai Jiang conceived the study and wrote the manuscript. Jiao Yu and Lin Shan performed COSMIC data analysis with help from Hongyu Ding, Mingxian Liu, Liang Xu, Jie Zhao, Zhengjin He, Shishuang Chen and Ailing Yang. Lin Shan performed studies of p53. Jiao Yu performed studies of PTEN and CDKN2A. Lin Shan and Jiao Yu prepared figures and analyzed data. All authors discussed the results and commented on the manuscript. Hai Jiang supervised the study.

## Acknowledgments

This work was supported by the major scientific research project (Grant No. 2017YFA0504503) from the Ministry of Science and Technology of China, “Strategic Priority Research Program” of the Chinese Academy of Sciences, Grant No. XDB19000000, and Natural Science Foundation of the People’s Republic of China (Grant No. 81572756). We thank Drs. Dangsheng Li, Zhaocai Zhou, Ning Gao, Jinqiu Zhou and Yun Zhao for discussions and helpful comments, and Animal Core Facility and Core Facility for Cell Biology at SIBCB.

## Supplementary information

**Figure S1.**
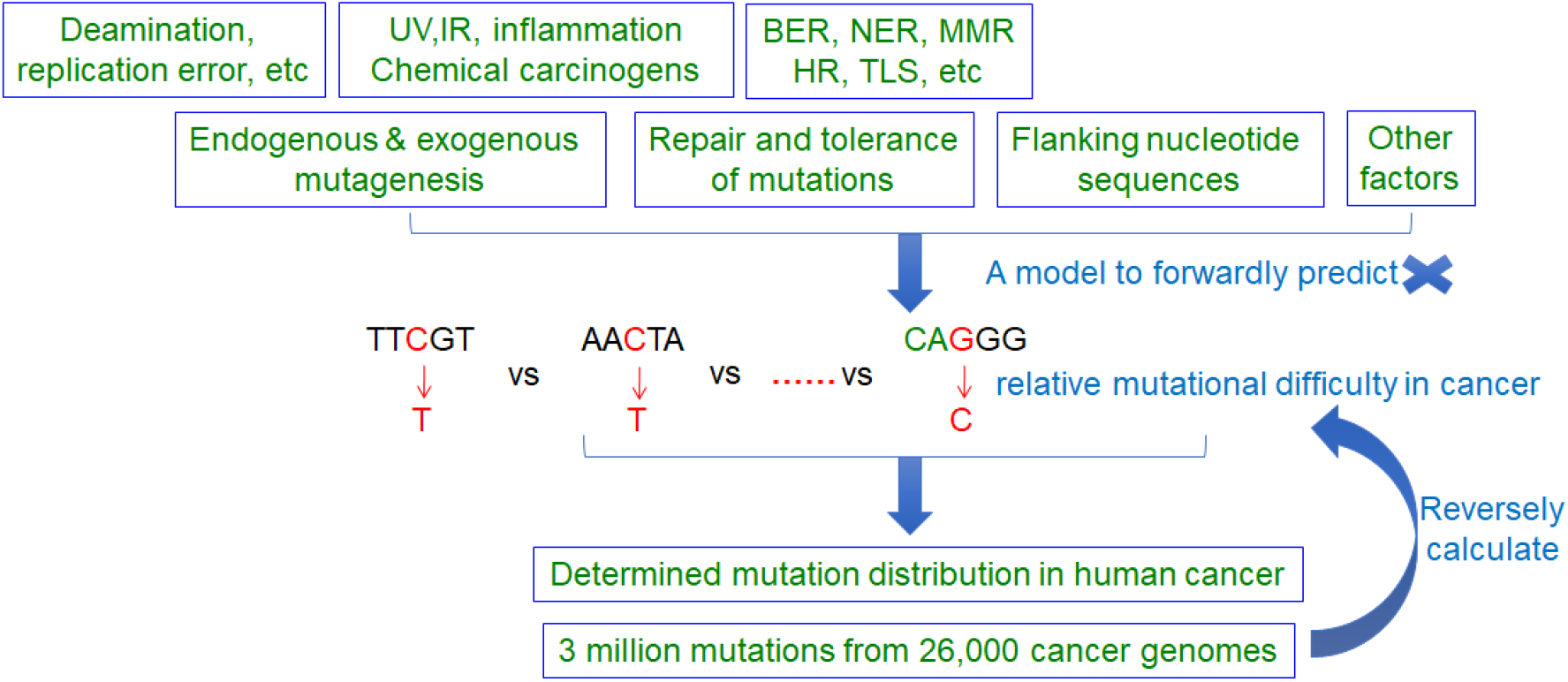
Rationale for reverse calculation of relative mutational difficulty in human cancer.

**Figure S2.**
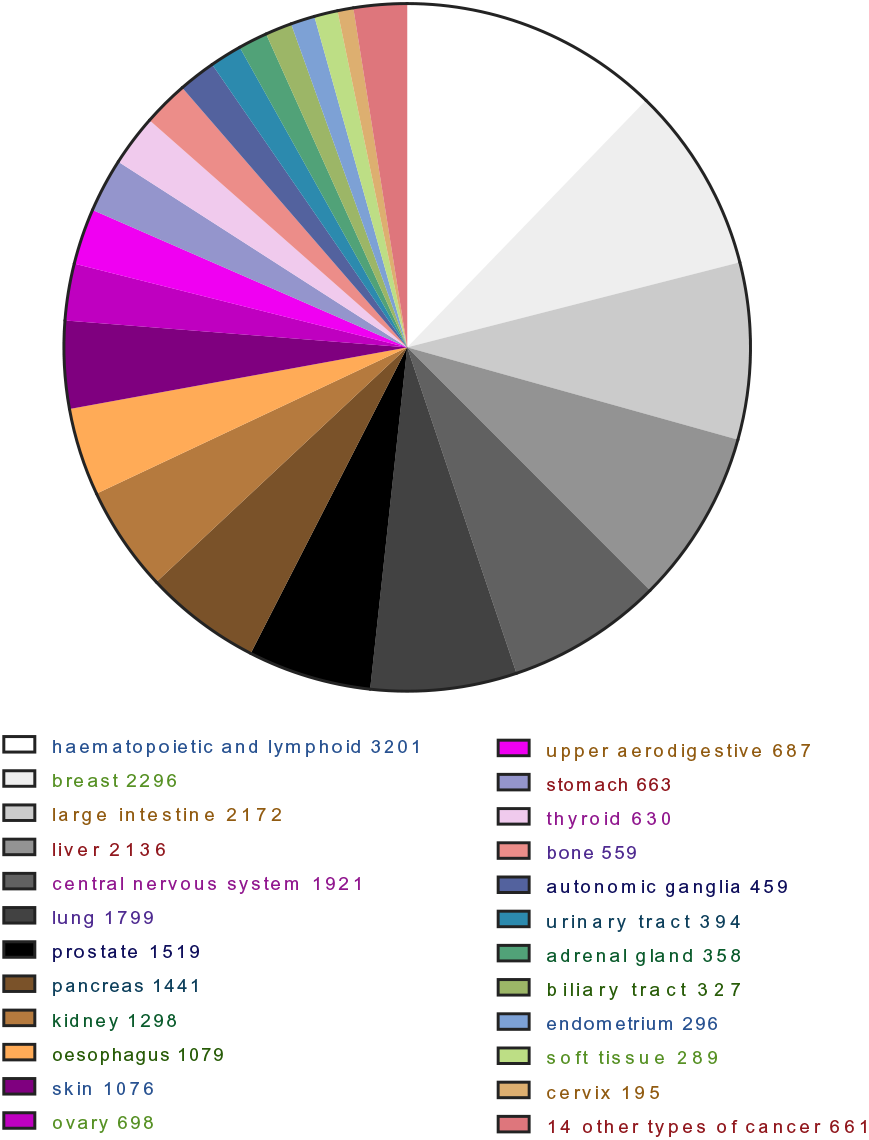
Composition of the 26,000 cancer samples with genomic sequencing data in COSMIC. The numbers following each cancer type indicate how many cancer genomes of the corresponding cancer type are analyzed.

**Figure S3.**
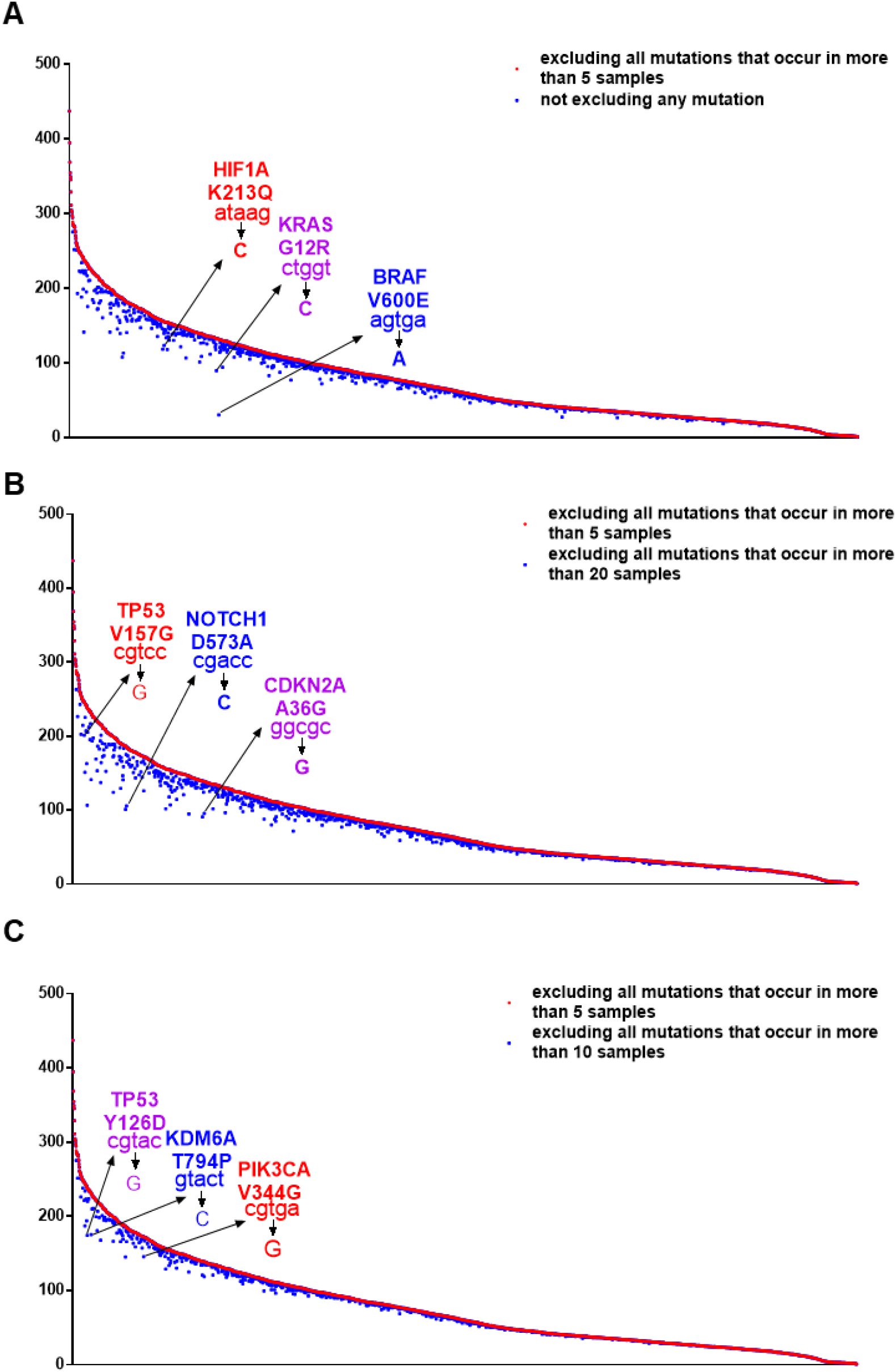
Potential impacts of highly-selected cancer mutations on calculation of relative mutational difficulty. Certain cancer-driving mutations such as BRAF V600E are highly selectively enriched in cancer. The number of such mutations are significantly increased in the dataset, not because they are easy to generate, but because they are strongly enriched during the tumorigenesis process. Therefore, their presence in the dataset may skew our estimation of the natural mutational tendency for each type of mutation. In our analysis, we excluded cancer mutations that occur in more than 5 cancer samples. The above panels show if no mutations are excluded (A), excluding mutations that occur in more than 20 (B) or 10 (C) samples, what is the impact on calculation of relative mutational difficulty. For example, if no mutations are excluded, the mutational difficulty for T to A substitution on a AGTGA sequence, which underlies the BRAF V600E mutation, will appear significantly lower. Several other similar examples are marked out in the above panels.

**Figure S4.**
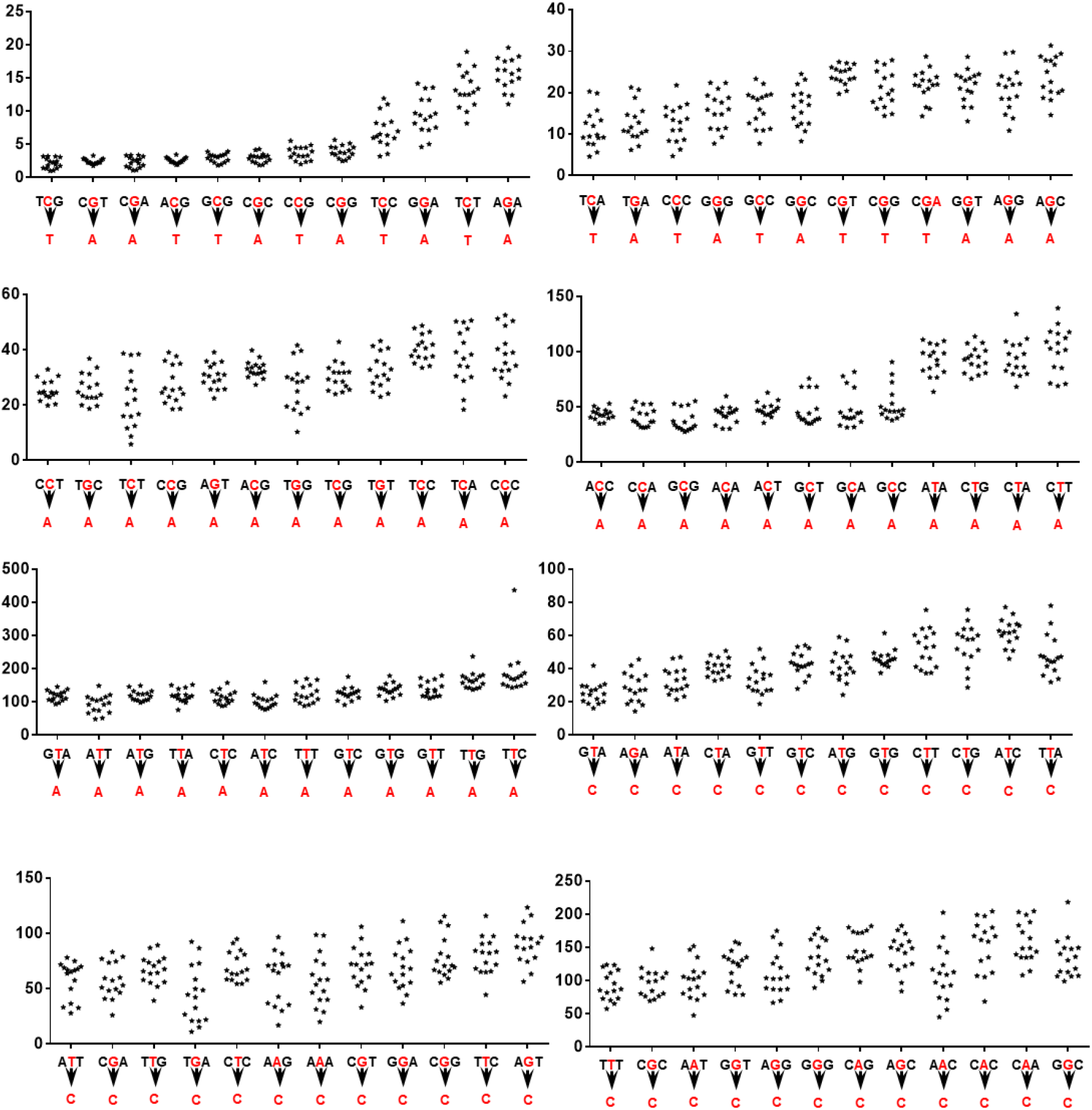
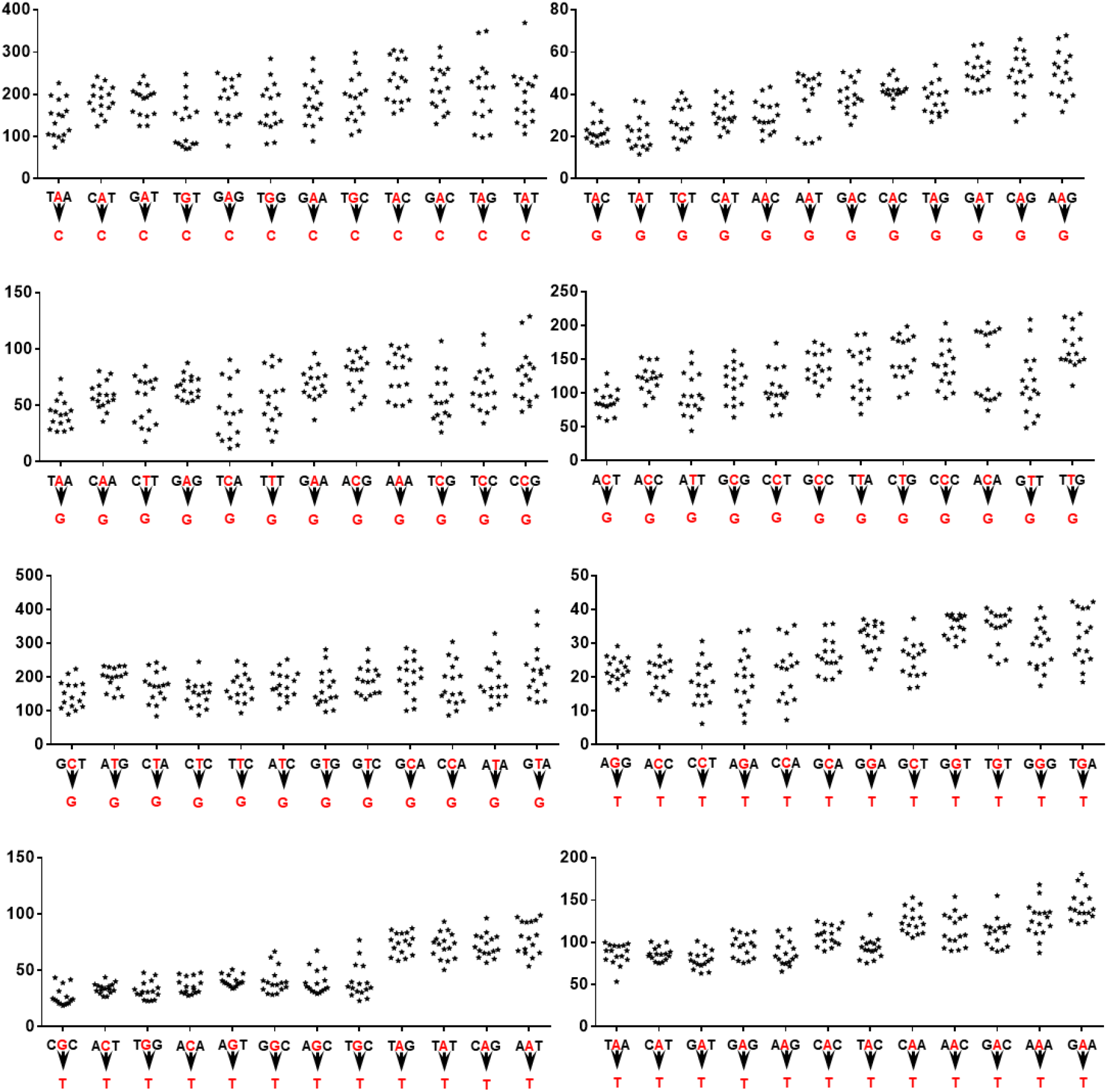
The influence of +2 and −2 nucleotides on relative mutational difficulty. In the above panels, the y-axis represents relative mutational difficulty. For example, in the last column of the lowest right panel, the sixteen * indicate on a NGAGN sequence, how 16 variations of the nucleotide in the +2 and −2 position will impact the relative mutational difficulty for A to T mutation on such NGAGN sequences.

**Figure S5.**
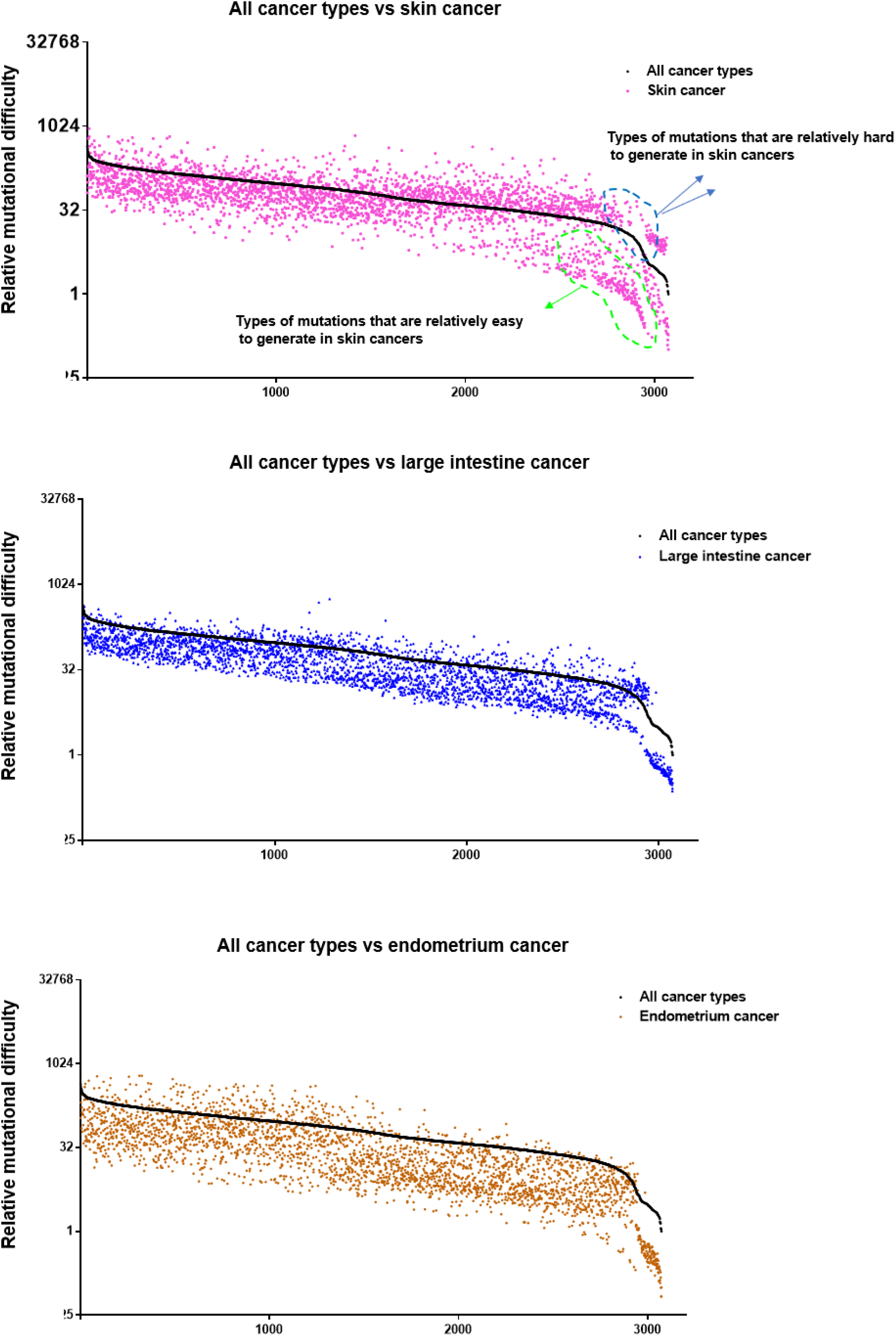
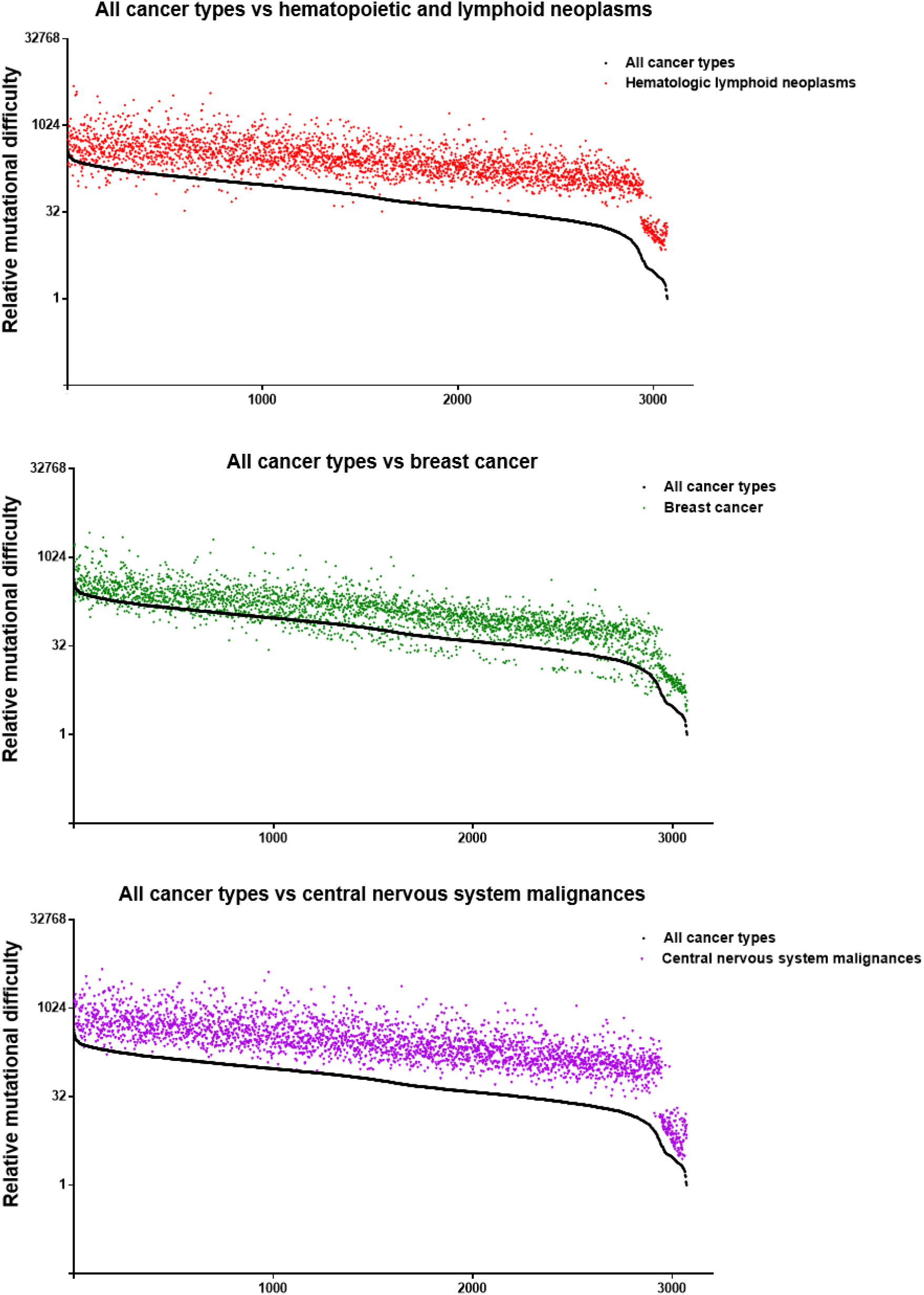
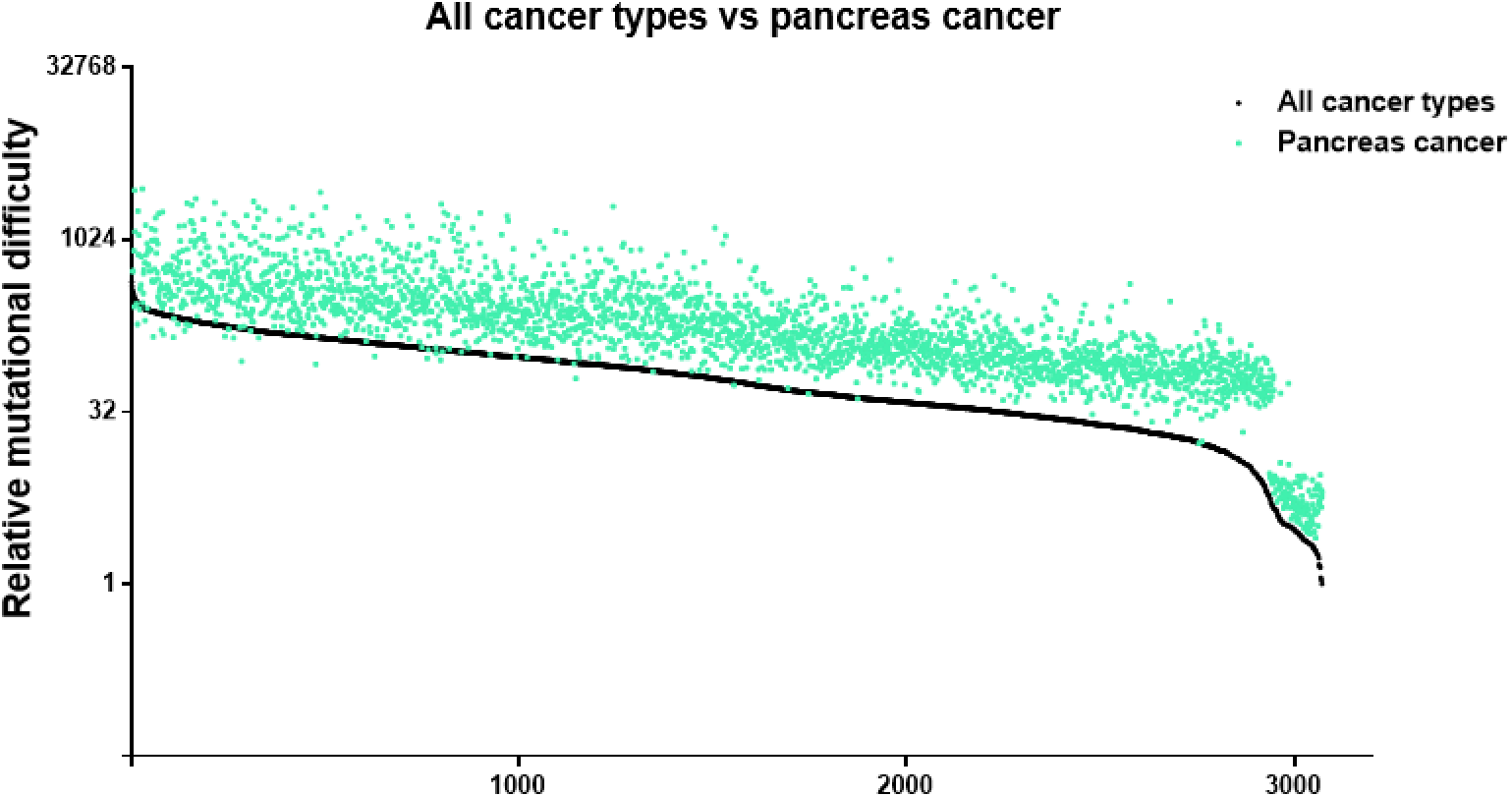
Cancer type-specific mutational difficulties compared with mutational difficulties calculated from all cancer types in average. Black dots indicate relative mutational difficulty scores calculated from 26,154 cancer genomes (all cancer types combined). In the top panel, examples of mutation types that are easier or more difficulty to generate in skin cancer are circled out.

**Figure S6.**
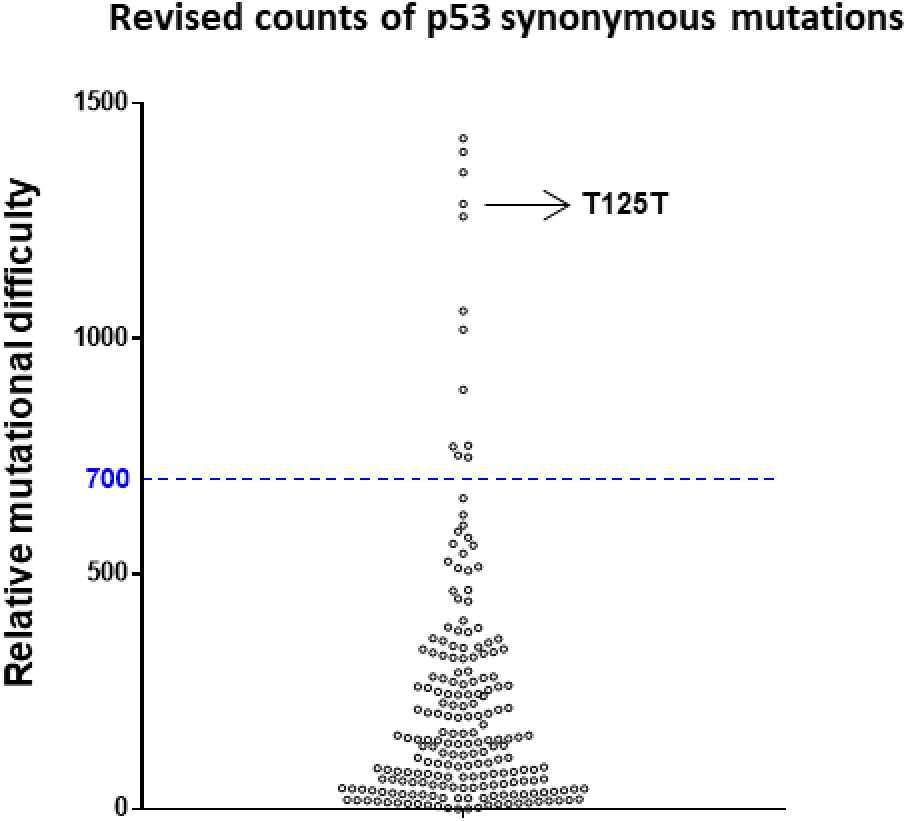
Distribution of revised counts of p53 synonymous mutations. Synonymous p53 mutations were compiled from the COSMIC database. Shown are revised mutation counts calculated for each type of synonymous p53 mutation. Certain synonymous mutations on p53 are known to be detrimental to the gene. For example, the T125T(c.375 G to A/C/T) mutation disrupts p53 splicing and causes loss of p53 activity(Supek et al., 2014).

**Figure S7.**
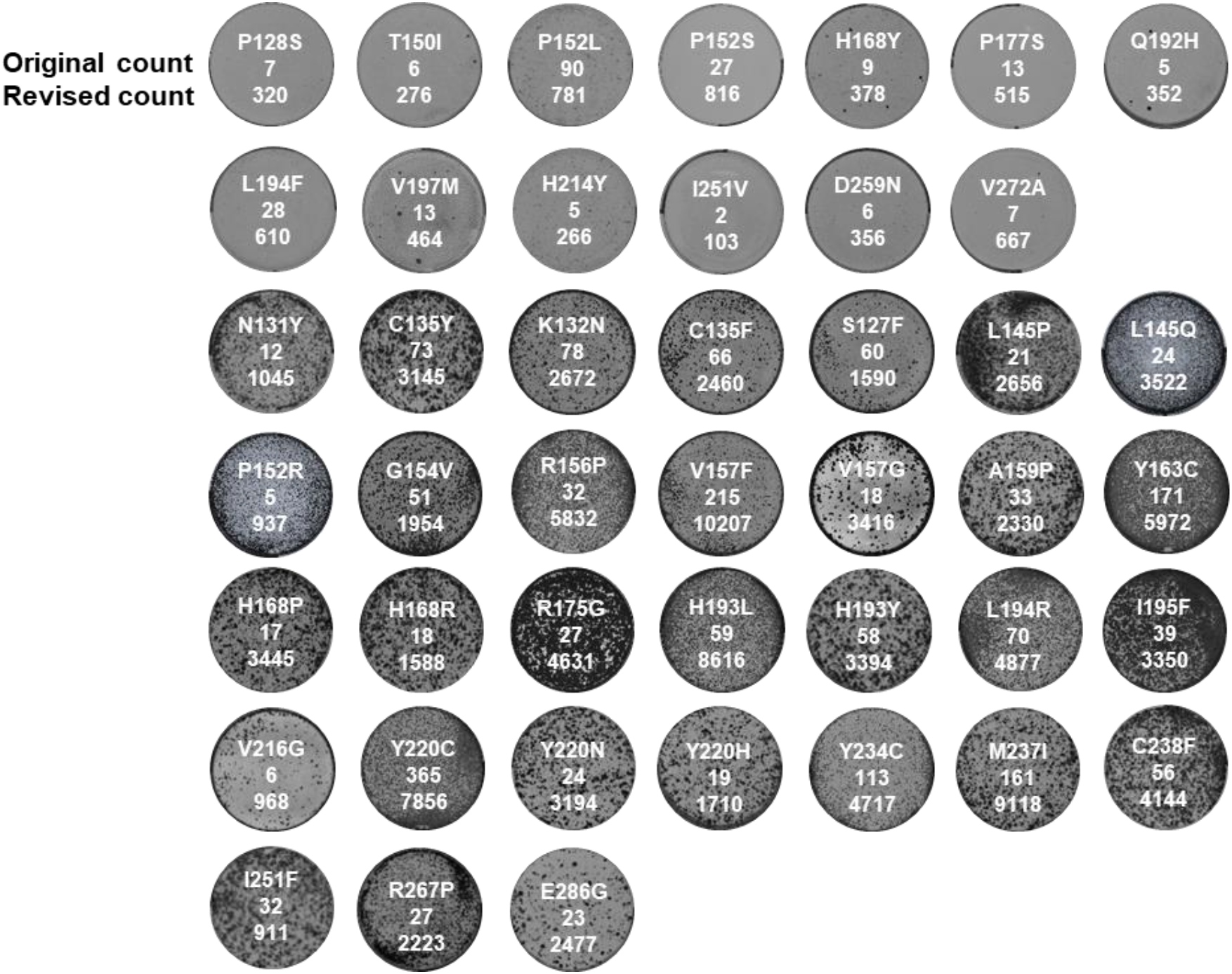
Saos-2 colony formation assay results for various p53 mutations. Shown here are experimental results of p53 mutations included in this study, in addition to those shown in Figure 2C, 2D and 4D. The original counts and revised counts are listed below each mutation.

**Figure S8.**
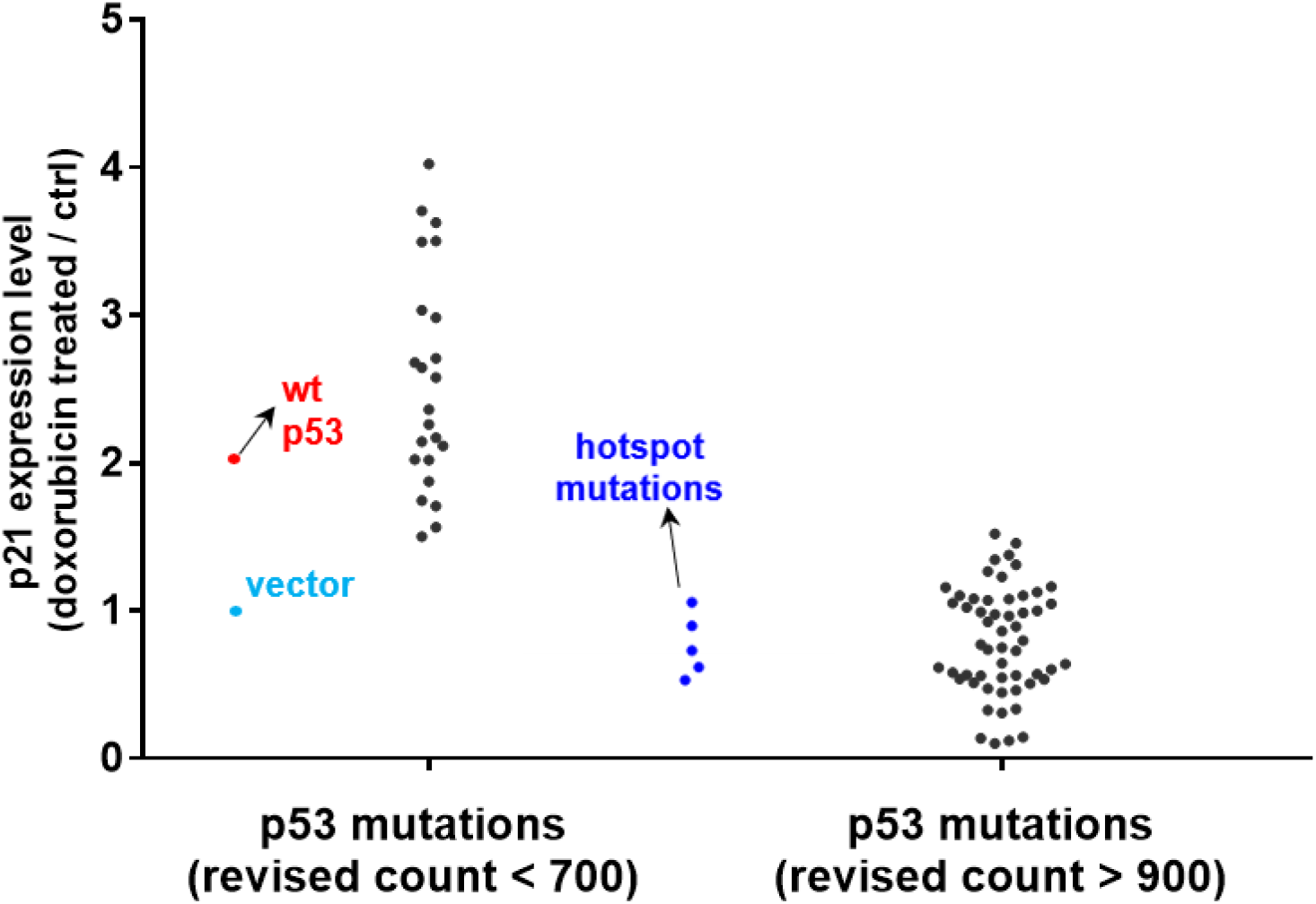
Transcriptional activities of different p53 mutants. Lentivirus encoding mutant or wild type p53 was used to infect HCT115 p53-/-cells at 30-40% infection rate. A puromycin selection marker in the lentivirus were used to select infected cells. Such cells were then treated with doxorubicin for 24 hours, and the transcriptional activity of p53 mutants were analyzed by extent of p21 upregulation.

**Figure S9.**
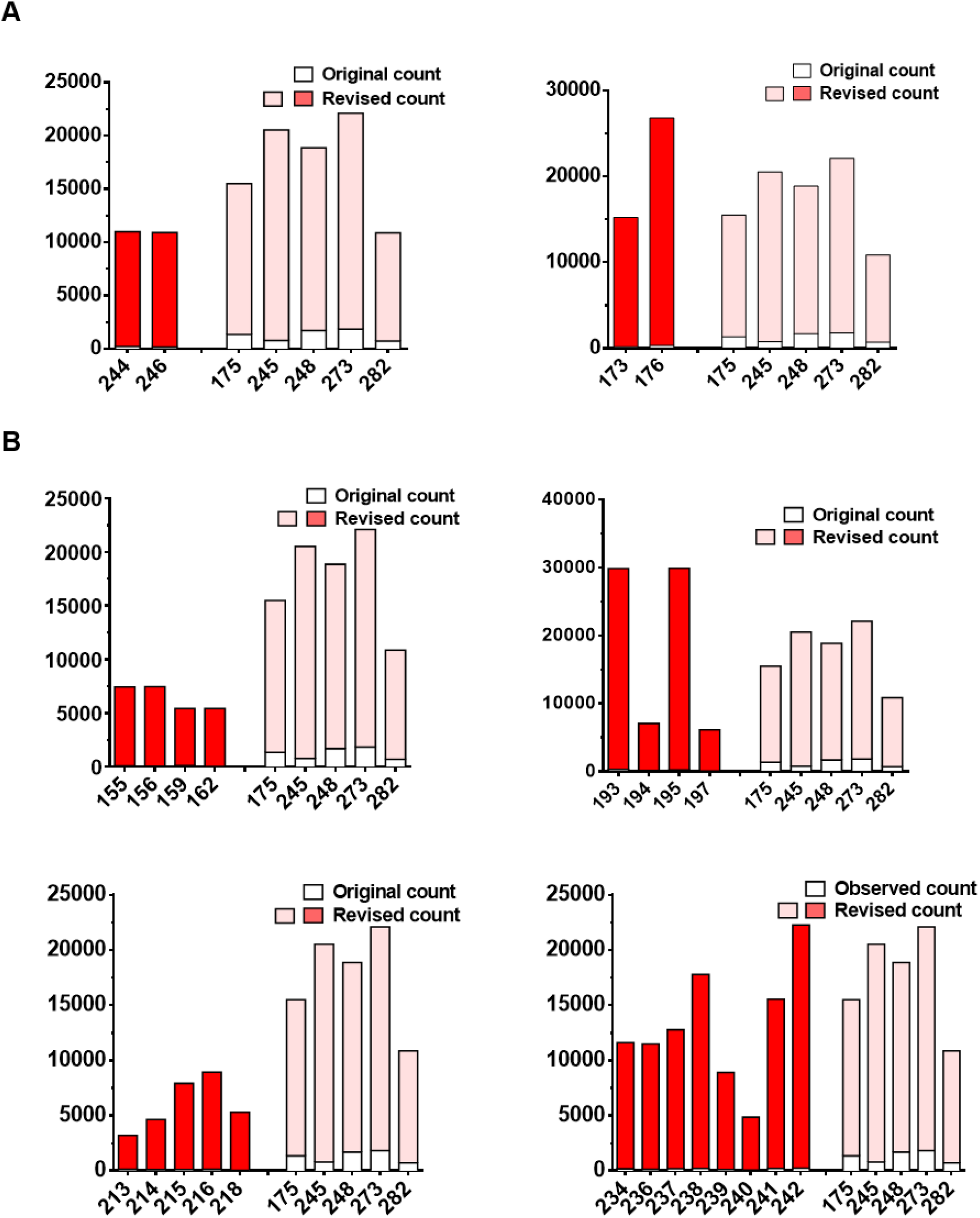
Additional functionally important amino acid residues and regions in p53. Original mutation counts of listed amino acid residues are indicated by white boxes, whereas revised counts are indicated by red or pink boxes. Hotspot mutation sites such as R175 and G245 are included as controls. Shown here are functionally important amino acids in addition to those shown in Figure 4.

**Figure S10.**
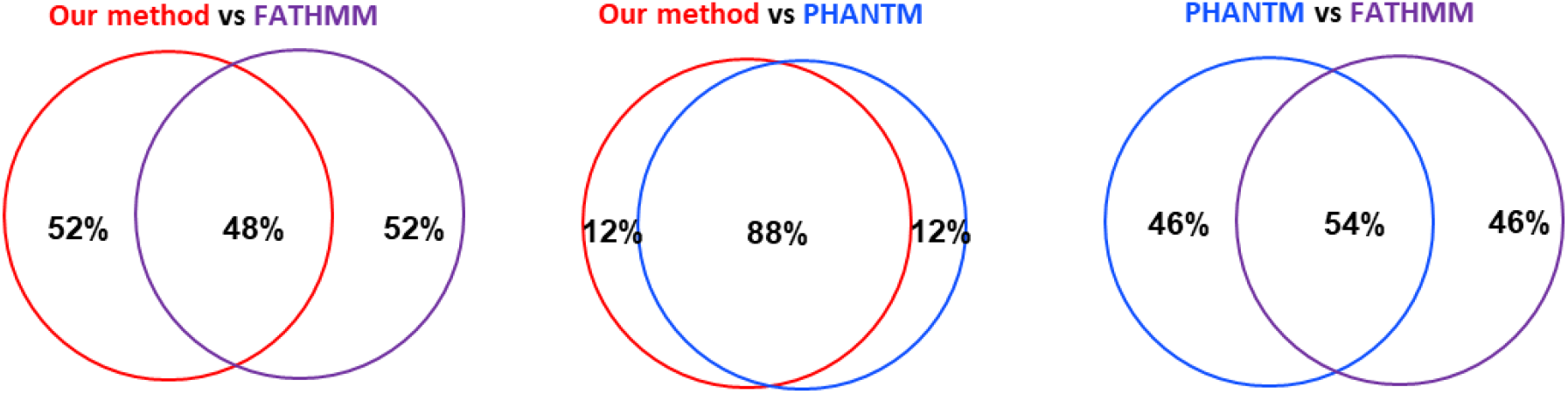
Comparison of functional predictions based on different methods. FATHMM annotation of p53 mutants are available at individual p53 mutation page on COSMIC site. 48% of what our method predicted to be wild type function or loss of function mutations are predicted in the same manner by the FATHMM method. PHANTM, Phenotypic Annotation of TP53 Mutations (http://mutantp53.broadinstitute.org/) provides functional assessment of individual p53 mutant, based on Giacomelli et al 2019. In that study, an extensive library of p53 mutants was introduced into cells, which were then treated under different conditions. The funtional status of p53 mutants was estimated based on whether they are tolerated by cells, as determined by massively parralle sequencing.Prediction by our method, which takes relative mutational difficulty into consideration, are highly consistent with the PHANTM results, with an 88% overlap. P53 mutations whose functional status are differently annotated between our method and FATHMM or PHANTM are listed in Table S4 and S5 respectively.

**Figure S11.**
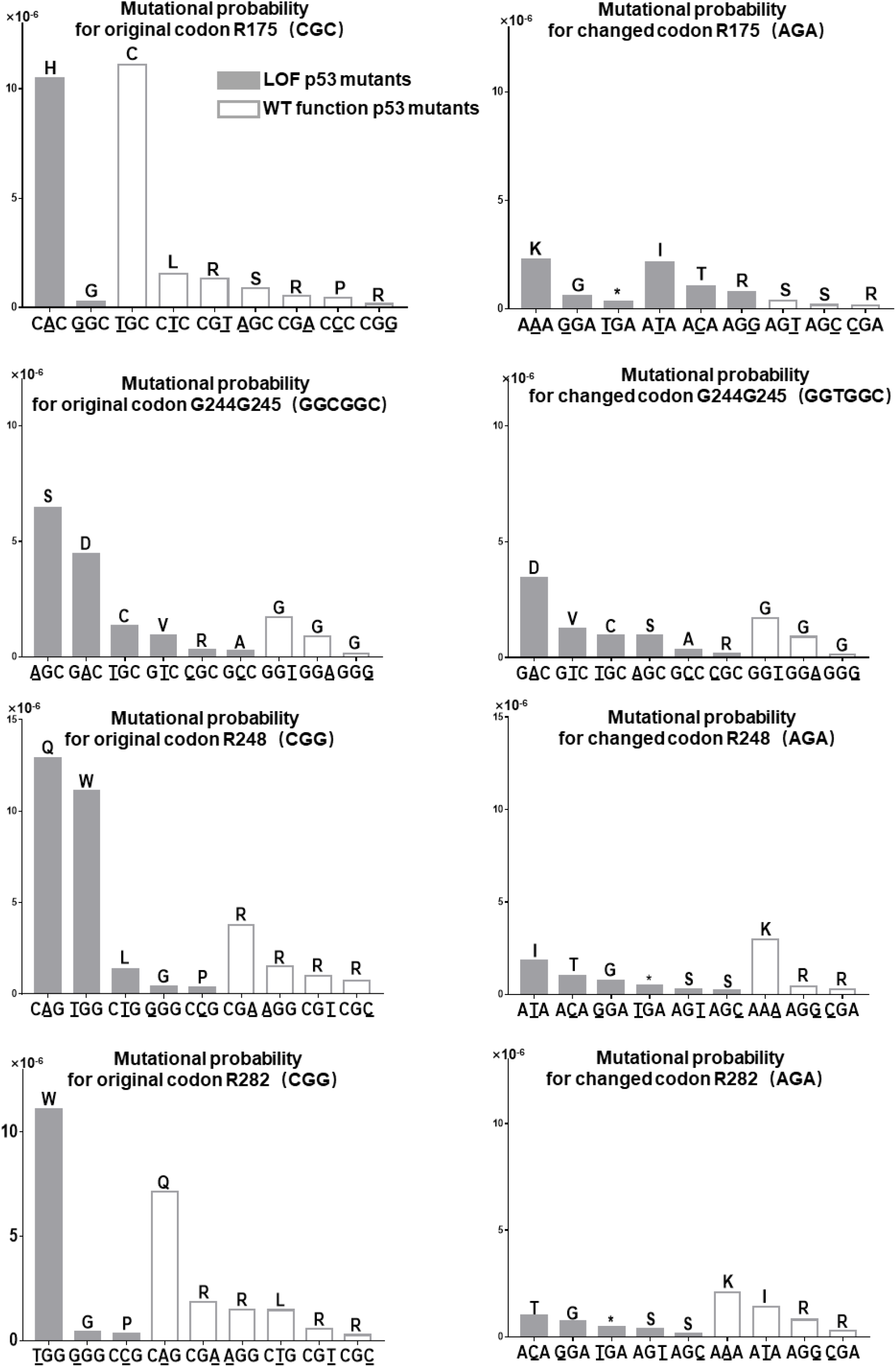
Mutational probability of the original and changed codons on p53 hotspot mutation sites. Original and changed codons for indicated p53 hotspot mutation sites are shown in these panels. Loss of function mutations are shown in dark grey. Mutations with wild type p53 function are shown in white. Shown here are analysis results for R175, G245, R248 and R282, and results for R273 is shown in Fig. 6B. Of note, the high mutation rate on G245 is enabled by the preceding codon G244, resulting in a nucleotide sequence (GC**<u>G</u>**<u>GC</u>) that is highly prone to **<u>G</u>** to A mutation, which causes the G245 (<u>GGC</u>) codon to be a hotspot mutation site. Change the G244 codon from GGC to GGT will decrease the mutation tendency of G245, without increasing mutation tendency of G244.

**Table S1** Calculation of relative mutational difficulty (All cancer types combined).

**Table S2** Cancer type-specific mutational difficulty.

**Table S3** Original and revised mutation counts of p53 mutations in COSMIC database.

**Table S4** List of p53 mutations whose functionalities are annotated differently by revised counts method and FATHMM.

**Table S5** List of p53 mutations whose functionalities are annotated differently by revised counts method and PHANTM.

**Table S6** List of p53, PTEN and INK4A mutants analyzed in this study.

**Table S7** List of primers used for mutagenesis experiments.

**Table S8** List of primers used for qPCR analysis.

